# Development of allosteric, selective cyclin-dependent kinase 2 (CDK2) inhibitors that are negatively cooperative with cyclin binding and show potential as contraceptive agents

**DOI:** 10.1101/2022.06.30.497818

**Authors:** Erik B. Faber, Jian Tang, Emily Roberts, Sornakala Ganeshkumar, Luxin Sun, Nan Wang, Damien Rasmussen, Abir Majumdar, Kristen John, An Yang, Hira Khalid, Jon E. Hawkinson, Nicholas M. Levinson, Ernst Schönbrunn, Vargheese Chennathukuzhi, Daniel A. Harki, Gunda I. Georg

**Affiliations:** Department of Medicinal Chemistry, University of Minnesota College of Pharmacy – Twin Cities, Minneapolis, MN; Institute for Therapeutics Discovery and Development, University of Minnesota College of Pharmacy – Twin Cities, Minneapolis, MN; Medical Scientist Training Program, University of Minnesota Medical School – Twin Cities, Minneapolis, MN; Department of Molecular and Integrative Physiology, University of Kansas Medical Center, Kansas City, KS; Drug Discovery Department, Moffitt Cancer Center, Tampa, FL; Department of Pharmacology, University of Minnesota Medical School – Twin Cities, Minneapolis, MN; Department of Biochemistry, Molecular Biology & Biophysics, University of Minnesota Medical School – Twin Cities, Minneapolis, MN

## Abstract

Compared to most ATP-site kinase inhibitors, small molecules that target an allosteric pocket have the potential for improved selectivity due to the often observed lower structural similarity at these distal sites. Despite their promise, relatively few examples of structurally confirmed, high-affinity allosteric kinase inhibitors exist. Cyclin-dependent kinase 2 (CDK2) is a target for many therapeutic indications, including non-hormonal contraception.^1^ However, an inhibitor against this kinase with exquisite selectivity has not reached the market because of the structural similarity between CDKs.^1-2^ In this paper, we describe the development and mechanism of action of new type III inhibitors that bind CDK2 with nanomolar affinity, making them the highest affinity, structurally confirmed allosteric CDK inhibitors reported. Notably, these anthranilic acid inhibitors exhibit a strong negative cooperative relationship with cyclin binding, which remains an underexplored mechanism for CDK2 inhibition. Furthermore, the binding profile of these compounds in both biophysical and cellular assays demonstrate the promise of this series for further development into a therapeutic selective for CDK2 over highly similar kinases like CDK1. The potential of these inhibitors as efficacious contraceptive agents is seen by incubation with mouse testicular explants, where they recapitulate *Cdk2*^*-/-*^ and *Spdya*^*-/-*^ phenotypes.

## Introduction

Kinases remain attractive therapeutic targets to treat a variety of diseases, but with over 500 kinases in the human proteome, the selective targeting of a particular kinase remains challenging. This is in large part due to the structural similarity among kinases, especially at the ATP-site where most kinase inhibitors bind. To meet this challenge, a variety of strategies have emerged including developing inhibitors that either bind unique inactive conformations, target unique residues including nucleophilic residues for covalent attachment,^3^ or are heterobifunctional and lead to kinase degradation.^4^ In addition, allosteric inhibitors, or compounds that bind sites on the kinase away from the ATP-site, have been a promising avenue to selectively inhibit kinases as these sites often have reduced structural similarity. Some allosteric kinase inhibitors even demonstrate selectivity for oncologic mutant isoforms of the same kinase.^5-6^ To date, four allosteric kinase inhibitors have been approved by the U.S. Food and Drug Administration to treat BRAF-mutated cancers and recently asciminib was approved as a treatment for chronic myeloid leukemia targeting an allosteric pocket of ABL kinase.^7-11^ Despite this success, few high affinity and well-characterized allosteric kinase inhibitors have been reported in the literature. In this paper, we will showcase a series of anthranilic acid-derived inhibitors that bind an allosteric pocket of cyclin-dependent kinase 2 (CDK2) with nanomolar affinity.

CDK2 is a member of the CDK family of kinases that control the cell cycle and is activated by binding E-type cyclins at the G1-S transition, then by A-type cyclins in S phase. Additionally, CDK2 must be phosphorylated on the activation loop to become fully activated. CDK2 also plays a critical role in meiosis since *Cdk2*^*-/-*^ mice are sterile but otherwise healthy,^12-13^ making CDK2 a validated target for non-hormonal contraceptive development.^1^ In meiosis, the protein Speedy1 (SPY1) binds unphosphorylated CDK2 at the same site where cyclins bind^14^ and appears to be the crucial binding partner of CDK2 in the pachytene stage of prophase I.^15^ While the *Cdk2*^*-/-*^ phenotype suggests a selective CDK2 inhibitor will have a favorable safety profile, further investigation reveals that selective inhibition of CDK2/cyclin complexes can be relatively toxic for healthy cells due to the sequestration of cyclins and thus the loss of compensatory kinase activation.^16^ However, a compound that disrupts the protein-protein interaction (PPI) between CDK2 and its protein partners like cyclin or SPY1 has the potential to be an alternative, less toxic CDK2 inhibitor. In this manner, an ideal inhibitor would free cyclins normally bound to CDK2, leading to A) the activation of compensatory kinases by displaced cyclins in somatic cells and B) disruption of the CDK2/SPY1 interaction crucial for meiosis in spermatocytes, providing a safe and efficacious contraceptive option.

We previously discovered that the dye 8-anilino-1-naphthalene sulfonic acid (**ANS**) bound an unrecognized allosteric pocket within CDK2 with moderate affinity with a 5-to 10-fold enhancement in affinity in the presence of certain orthosteric inhibitors.^17-18^ Despite this promise, **ANS** is known to be a promiscuous protein binder^19-23^ and contains a sulfonic acid moiety that confers poor pharmacokinetic properties. However, we exploited the environmentally sensitive fluorescent nature of **ANS** to develop a high-throughput assay and used it to discover new chemical scaffolds with improved physicochemical properties that bind the allosteric pocket with higher affinity.^24^ From those efforts, we herein reveal the highest affinity allosteric CDK inhibitors reported. We confirm their type III allosteric interaction by X-ray crystallography and use other biophysical techniques to demonstrate negative cooperativity with cyclin binding. This cooperative relationship continues to hold promise as an underexplored mechanism of CDK2 inhibition, only observed with **ANS** prior.^17^ Furthermore, the challenge of developing selectivity for CDK2 over structurally similar kinases like CDK1 has been overcome, as seen in both biophysical and cellular assays. Finally, we show that mouse testes incubated with our allosteric CDK2 inhibitors recapitulate features of a *Cdk2*^*-/-*^ phenotype.

## Results

We discovered that anthranilic acids bind the **ANS** allosteric pocket of CDK2. The first compound crystallized of this series was **1**, which binds CDK2 with low micromolar affinity as measured by isothermal titration calorimetry (ITC) and by surface plasmon resonance (SPR) using the steady-state approximation (SSA) affinity fit (Fig. 1, SI Fig. 1).

**Fig. 1.**
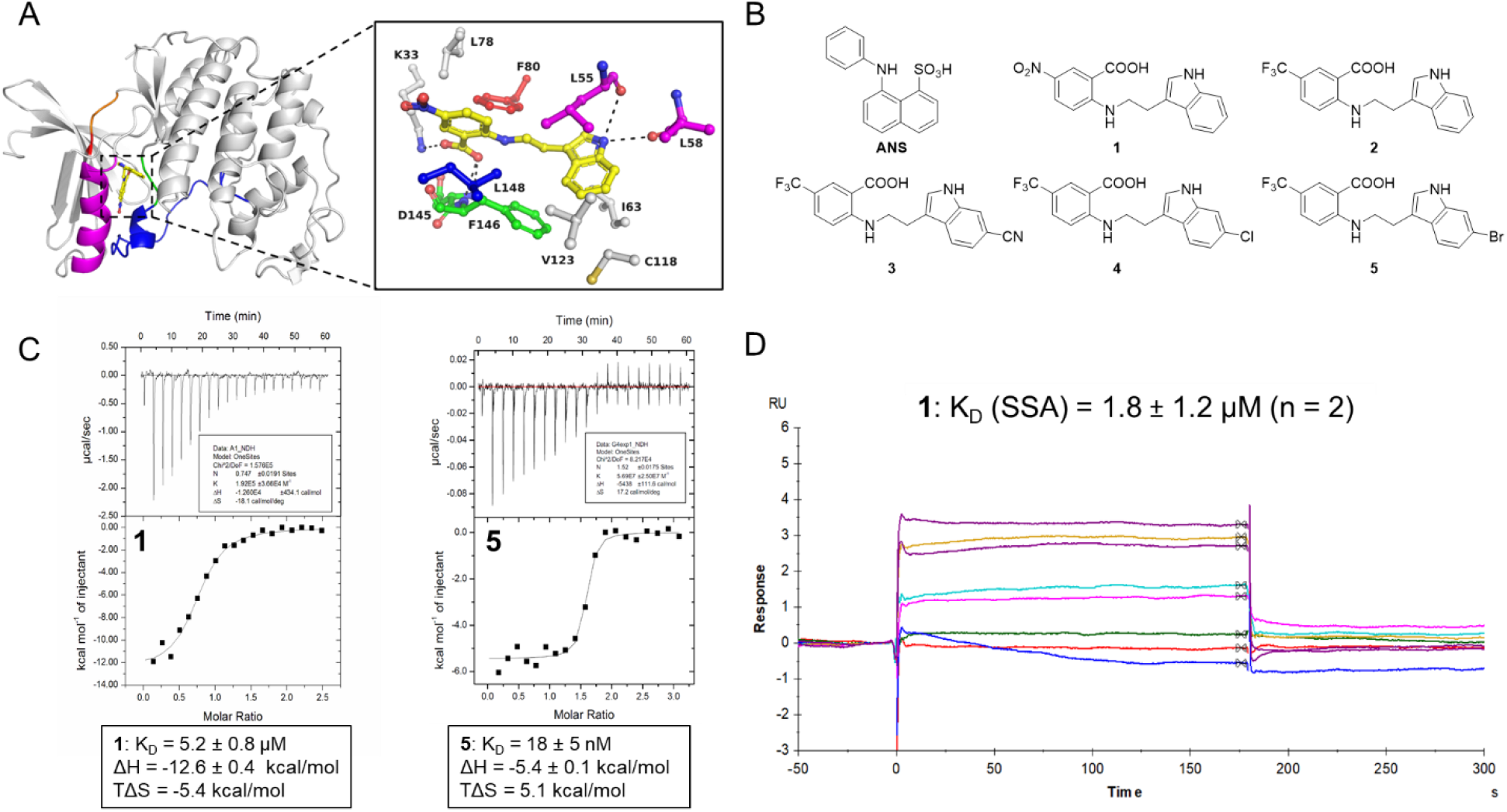
Binding characterization of anthranilic acids as CDK2 allosteric inhibitors. A. Cocrystal structure of **1** (yellow) bound to an allosteric pocket of CDK2 (PDB ID 7RWF). The C-helix is shown in magenta, the activation loop in blue, the DFG motif in green, the hinge region in orange, and the gatekeeper residue in red. H-bonds are indicated as black dotted lines (hydrophobic van der Waals interactions and water molecules were omitted for clarity). B. Chemical structures of CDK2 allosteric inhibitors. C. ITC traces and thermodynamic binding values of **1** (left) and **5** (right). D. Representative SPR trace and KD of **1** against CDK2, both replicates with individual KD values (using the steady-state approximation, SSA) from each run in SI Fig. 1.

The carboxylic acid of **1** forms a charge-charge interaction with the side chain of the catalytic residue K33 in the hydrophobic interior of the protein, the same residue that engages the sulfonic acid in **ANS** (Fig. 1A). Due to the strength of this type of interaction in a nonpolar environment, it is likely crucial for the binding of these allosteric inhibitors. Additionally, F80 (gatekeeper residue of ATP site) and F146 (of the DFG motif) engage in π-π interactions with the benzene and indole rings of **1**, respectively. The N-H of the indole forms hydrogen bond interactions with the backbone carbonyls of L55 and L58 of the C-helix, and the remainder of the binding interaction seem to be driven by van der Waals forces in a pocket largely composed of hydrophobic residues. Both **1** and **ANS** are amphipathic in nature, with a negatively charged moiety driving the binding to K33 and much of the remaining structure engaging with the hydrophobic pocket.

We next made two substantial improvements to the anthranilic acid series. First, we sought to replace the nitro group on the benzene ring, as this group can be toxic in biological systems.^25^ We find the trifluoromethyl group in **2** not only improves the affinity 2-fold but retains a strong enthalpic contribution of binding with a lesser entropic penalty (Fig. 1C, Table 1, SI Fig. 2). The binding pose of **2** to CDK2 is nearly identical to that of **1** (PDB ID 7S84). Next, substitutions at the 6 position of the indole improve the affinity substantially. Linear groups like nitrile (**3**) or larger halogens like chlorine (**4**) or bromine (**5**) at this position afford binding with less enthalpic benefit and greater entropic contribution (Fig. 1C, Table 1, SI Fig. 2). The nitrile group at this position on the indole in **3** may linearize the molecule compared to **2** and subsequently increase its solvation shell, which is broken when bound into the allosteric pocket. The increased hydrophobicity of **4** and **5** take this principle further, as binding of these compounds into the hydrophobic allosteric pocket led to a greater entropy increase. We were able to orthogonally confirm the affinities of many of these compounds with nanomolar affinities by SPR (SI Fig. 3).

**Table 1.**
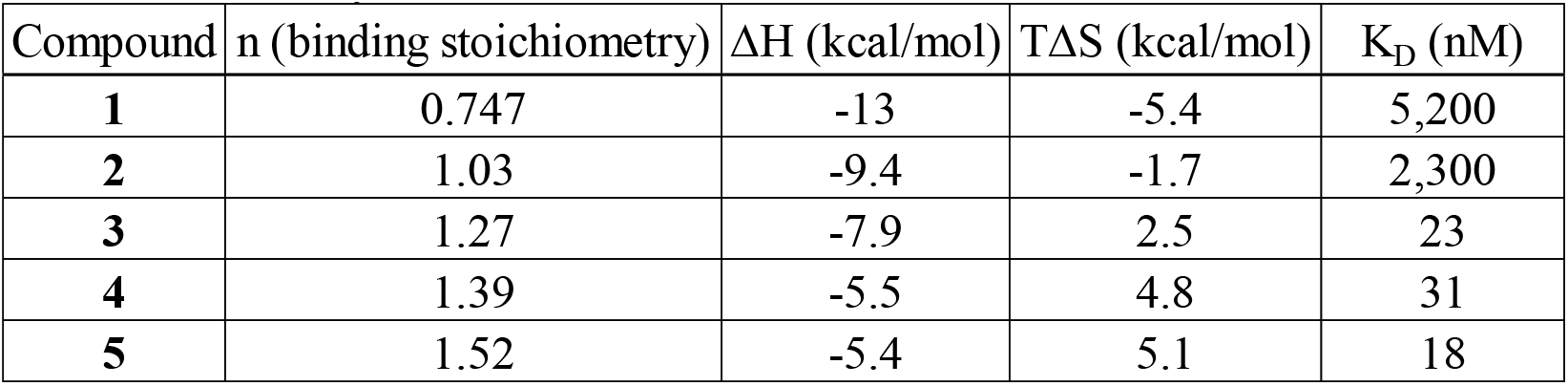
Enthalpic and entropic contributions of binding by each ligand and their respective affinity, as determined by ITC.

To explore the inhibitory potential of these compounds, **4** and **5** were selected for further biophysical testing. Despite its high affinity for CDK2, **4** poorly inhibits a variety of CDK/cyclin complexes in commercial kinase activity assays (SI Fig. 4). Further analysis of the crystal structure of **1** bound to CDK2 reveals that the allosteric inhibitor engages with L53 and L58 of the C-helix, potentially in a manner incompatible with cyclin A binding that would require a reorientation of the C-helix (Fig. 1A). It is likely that in this CDK panel, in which tight-binding cyclin is preincubated with each respective CDK enzyme, the binding of **4** is precluded by the CDK/cyclin PPI (i.e., the CDK2/cyclin A/A1/E/O complexes).

To test this hypothesis, we took advantage of an intramolecular Förster Resonance Energy Transfer (FRET) assay developed to indirectly track the position of the activation loop on CDK2 and thus various conformational states of the kinase. In this assay, CDK2 is modified by attaching an Alexa Fluor® donor/acceptor dye pair split between a relatively stable region of the kinase and the mobile activation loop.^26^ As cyclin A is titrated into CDK2 and the activation loop is stabilized in its active outward conformation, the donor:acceptor ratio of the dyes increases (Fig. 2A). In the absence of cyclin, when **4** and **5** are titrated into this CDK2 system, the donor:acceptor ratio shifts to a lower value, representing a different ensemble of inactive state(s) of CDK2 with corresponding K_D_ values of 280 nM and 200 nM, respectively (Fig. 2B, SI Fig. 5). However, as the cyclin A concentration is increased, the affinities of both small molecules decrease. For example, a cyclin A concentration of 1.9 μM increases the K_D_ of compound **5** to 1.5 μM, a change of nearly 10-fold (Fig. 2B). This change in affinity is even more pronounced for compound **4** (SI Fig. 5) and supports the negative cooperative relationship with cyclin binding that was hypothesized by structural analysis.

**Fig. 2.**
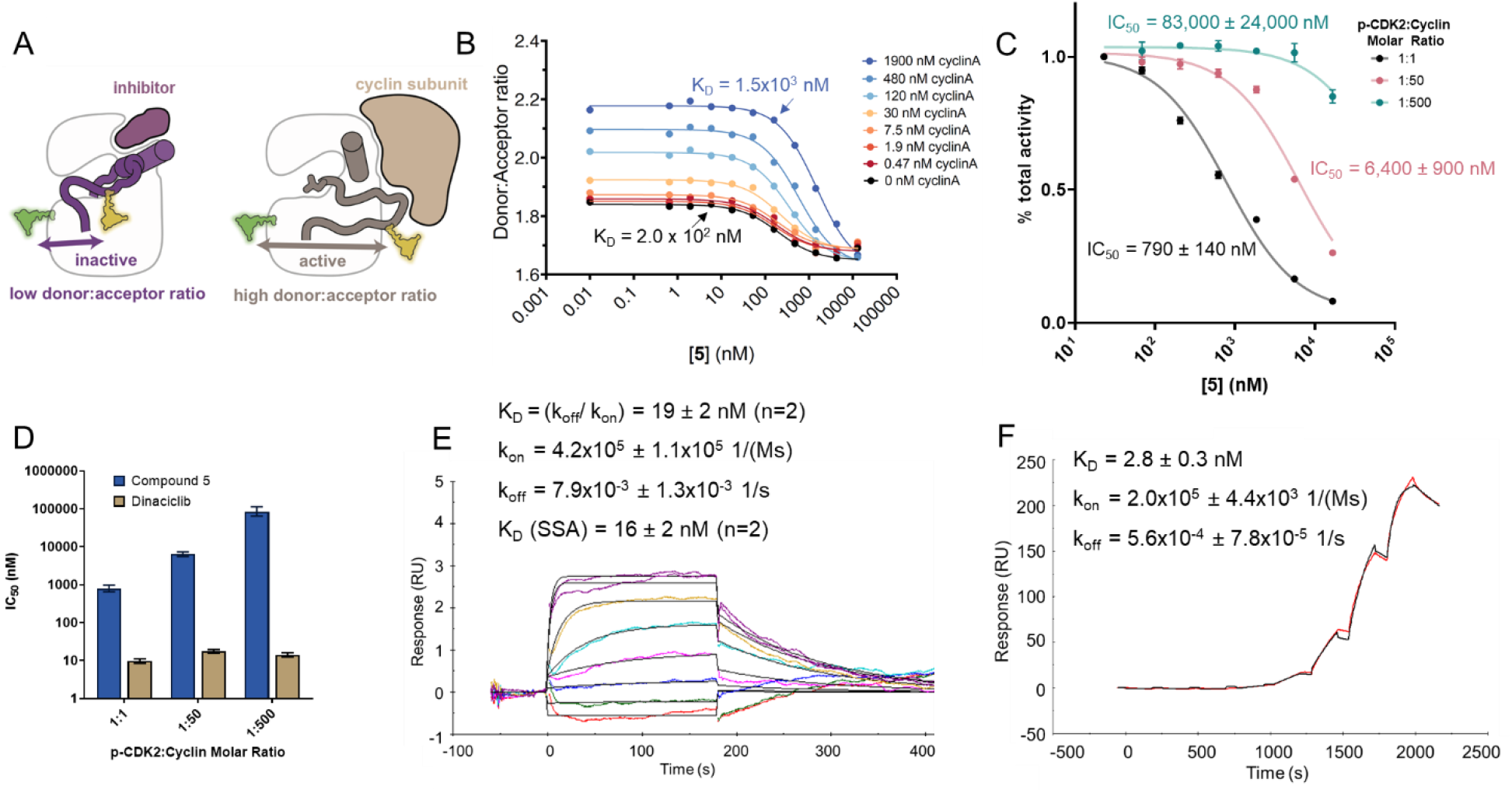
Compound 5 demonstrates negative cooperativity with cyclin A. A. An intramolecular FRET assay was developed for CDK2 to track the position of the activation loop. B. Compound **5** has lower affinity for CDK2 in the FRET assay as cyclin concentration increases. C. As cyclin concentration increases in an activity assay, the inhibitory potential of **5** decreases. D. The IC50 values of compound **5** compared to ATP-site inhibitor **dinaciclib** with varying p-CDK2:cyclin ratios. E. SPR data of **5** binding CDK2 reveals a fast on, fast off binding profile. F. SPR binding data of cyclin towards CDK2 reveals a slower off rate than that observed for small molecule inhibitors.

Next, we wanted to examine if this negative cooperativity is also observed in a CDK2 activity assay. We tested **5** rather than **4** due to its stronger affinity observed in the FRET experiments in the presence of cyclin A. Because traditional activity assays where the CDK2/cyclin complex is preformed do not demonstrate inhibition with our allosteric CDK2 inhibitors, we preincubated p-CDK2 with our allosteric inhibitor before adding varying amounts of cyclin A. At a 1:1 p-CDK2:cyclin molar ratio, the IC_50_ of **5** is ∼8-fold more potent than at a 1:50 ratio and >100-fold more potent than at a 1:500 ratio, showing a negative cooperative relationship with cyclin (Fig. 2C-D). The negative cooperativity observed in this kinase activity assay agrees with the conformational FRET assay (Fig. 2B). In contrast, the potency of ATP-site inhibitor **dinaciclib** is largely independent of cyclin concentration (Fig. 2D, SI Fig. 6).

To understand why the order of allosteric inhibitor and cyclin addition is important, we returned to the SPR assay to compare the binding kinetics of both our inhibitors and cyclin towards CDK2. As seen with previous CDK2 inhibitors, our allosteric inhibitors exhibit a fast association and fast dissociation rate (Fig. 2E, SI Fig. 1, SI Fig. 3).^2^ In contrast, cyclin has a similarly high affinity (K_D_ = 2.8 ± 0.3 nM, n = 2) but also has a 10+-fold slower dissociation rate compared to the allosteric inhibitors tested (Fig. 2E-F, SI Fig. 3, SI Fig. 7). Therefore, when cyclin A binds to CDK2, its slow dissociation precludes CDK2 from sampling the inactive conformation necessary for our allosteric inhibitors to bind. In contrast, the fast dissociation of the allosteric inhibitors allows cyclin to bind monomeric CDK2 and trap it in an active conformation, which is more pronounced as cyclin concentrations increase.

To explore the selectivity of these compounds, we next assessed target engagement in a cellular thermal shift assay (CETSA).^27^ Jurkat cells were selected for CETSA due to prior work showing a predictable T_agg_ across multiple, independent DMSO control experiments for CDK1 and CDK2 in this cell line.^28^ Additionally, Jurkat cells do not express a high amount of A- and E-type cyclins (Human Protein Atlas proteinatlas.org),^29^ making it possible to observe binding of our compounds to free, noncomplexed CDK2. Compound **5** is cell permeable, as shown by the intracellular engagement and thermal stabilization of CDK2 (ΔT_agg_ = 8.3 °C). No stabilization of CDK1 is observed (ΔT_agg_ = 0.4 °C), suggesting that **5** is selective for CDK2 over the very similar CDK1 in this cellular context (Fig. 3A, SI Fig. 8). However, the novelty of this selectivity is difficult to compare to previously developed ATP-site inhibitors, as nonselective type I CDK2 inhibitors were unsurprisingly cytotoxic in this assay (data not shown).

**Fig. 3.**
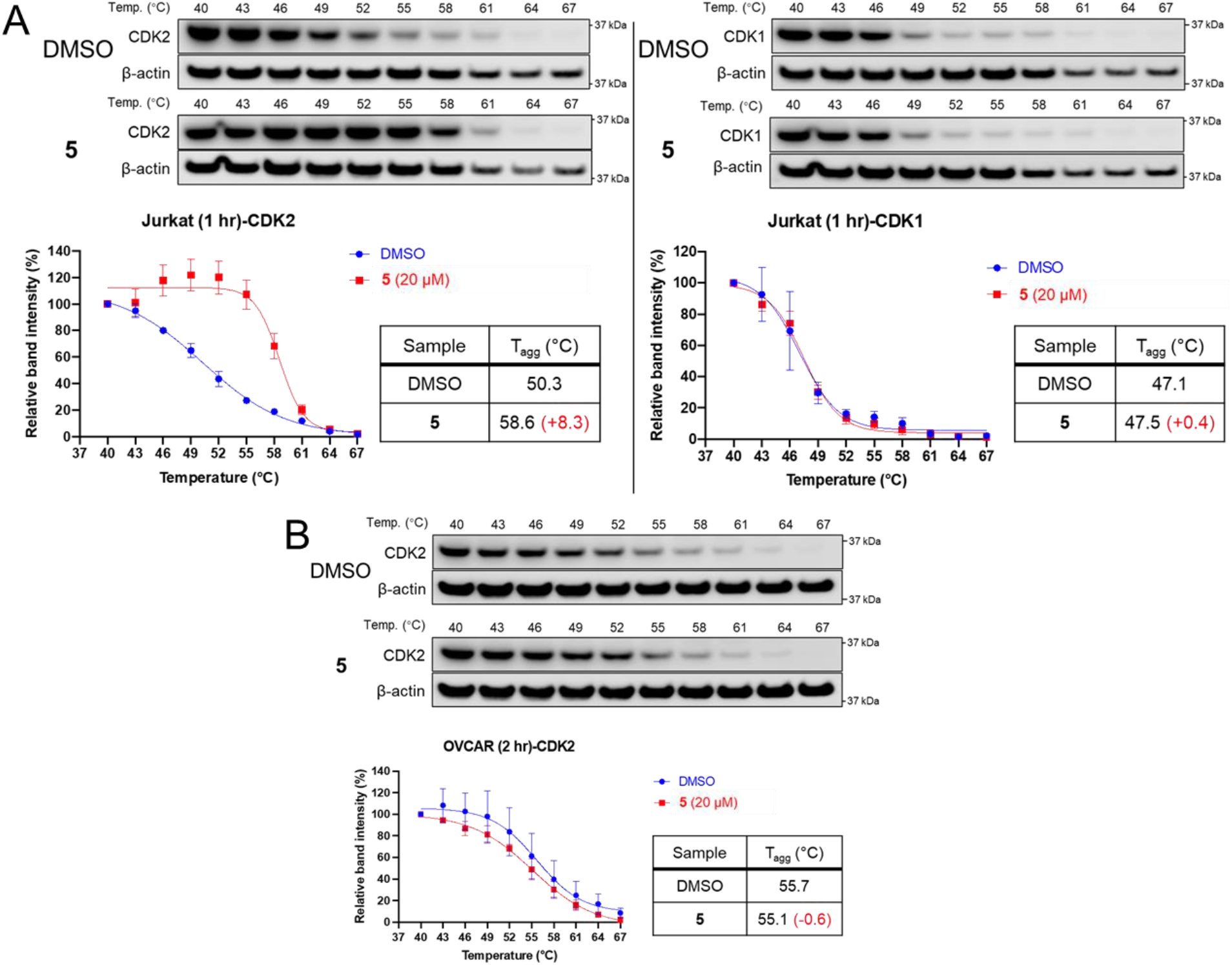
Target engagement of compound 5 as assessed by CETSA. A. Compound **5** stabilizes CDK2 over CDK1 intracellularly in CETSA experiments using Jurkat cells. B. Compound **5** does not stabilize CDK2 over DMSO control in CETSA experiments using OVCAR-3 cells. Protein abundance curves from the CETSA experiments indicate mean ± SEM from three biological replicates.

We sought to confirm the selectivity observed in the CETSA more directly in an orthogonal assay. Previous examination of ATP-site inhibitor **dinaciclib** under similar SPR conditions revealed a K_D_ (SSA) = 78 ± 16 nM for CDK2 and a K_D_ (SSA) = 1.8 ± 0.2 μM for CDK1.^2^ In contrast, we find compound **5** does not bind CDK1 while it binds CDK2 with a K_D_ (SSA) = 16 ± 2 nM (Fig. 2E, SI Fig. 3, SI Fig. 9). We confirmed the validity of our CDK1 protein by testing **dinaciclib**, which binds CDK1 similarly to what was discovered previously with a K_D_ (SSA) = 1.5 ± 0.4 μM (SI Fig. 9). Therefore, both the CETSA and SPR data suggest that allosteric inhibitor **5** has exquisite selectivity for CDK2 over CDK1. This selectivity is significant, as many other previously developed and clinically tested ATP-site CDK2 inhibitors like **dinaciclib** demonstrate off-target toxicity due to their inability to differentiate between these two kinases.^2, 30-31^ With this promising feature of our allosteric inhibitors, we next sought to confirm if the negative cooperativity between **5** and cyclin previously observed in the biophysical assays also translates to a cellular context.

To test this, we examined the target engagement of **5** in cancer cells driven by cyclin overexpression. We chose the ovarian cancer cell line OVCAR-3, where prior work has established via *in vitro* and *in vivo* genetic knockdown experiments that growth of OVCAR-3 cells is driven by cyclin E1 overexpression and dependent on CDK2. Additionally, as a demonstration of the dependence on CDK2, **dinaciclib** has previously shown significant toxicity in OVCAR-3 cells.^32^ Despite binding with CDK2 in Jurkat cells, **5** is relatively nontoxic in OVCAR-3 cells driven by cyclin E1 overexpression, in contrast to the positive control **staurosporine** (SI Fig. 10). To examine engagement with CDK2 in a high cyclin context, we performed similar CETSA experiments in OVCAR-3 cells as was initially done in Jurkat cells. These experiments show that in contrast to the profound stabilization **5** affords CDK2 in Jurkat cells, there is no stabilization of CDK2 by **5** in OVCAR-3 cells (Fig. 3B, SI Fig. 11). This is likely due to the high cyclin content in OVCAR-3 cells, which precludes binding of our compound to the allosteric pocket. Coupling the cellular CETSA data with the negative cooperativity with cyclin A observed in the structural FRET and kinase activity assays, **5** substantially binds CDK2 when cyclins are not overexpressed and can disrupt the interaction between CDK2 and its protein binding partners in these contexts.

To assess the utility of our compounds to disrupt CDK2 protein engagement in non-cyclin overexpressing contexts, their contraceptive potential was next explored. CDK2’s meiotic function is controlled via a cyclin-independent mechanism, namely the binding of a distinct activator protein, SPY1.^15^ In particular, the CDK2/SPY1 interaction is crucial for homologous chromosomal pairing and synapse formation, double-strand break processing, and facilitating crossing-over in prophase I; both *Cdk2*^-/-^ and *Spdya*^*-/-*^ spermatocytes exhibit a disruption of these events.^13, 15, 33^ Because the crystal structure of SPY1 bound to CDK2 reveals a smaller protein with fewer protein-protein contacts than cyclin proteins, it is feasible that the affinity of this interaction could be weaker than that between CDK2 and cyclin.^14^ This would lead to an inhibitor that selectively disrupts CDK2/SPY1 complexes but leaves CDK2/cyclin complexes intact. To assess the contraceptive potential of our compounds, we tested **5** in spermatocytes from mouse testicular explants, as we knew it was cell permeable and binds to CDK2 when cyclins are not overexpressed. After addition of **5**, we examined the chromosomal arrangement by staining for SYCP3 as well as the localization of RAD51 and CDK2. Normally, RAD51 localizes to synapses only transiently and therefore is captured by microscopy in asynapsed regions. In contrast, both *Cdk2*^*-/-*^ and *Spdya*^*-/-*^ spermatocytes arrest meiosis in such a unique fashion that RAD51 can be found localized to synapsed regions.^15^ Mouse testicular explants incubated with **5** show significantly more abnormal RAD51 staining at or near the telomeres in synapsed regions compared to control treated explants, indicating arrested and unrepaired DNA double strand breaks (Fig. 4A-B, *p* = 0.017). Additionally, more meiotic cells show abnormal CDK2 staining on chromosomes within the nuclei in the presence of **5** (50% of samples) compared to control samples (17%), indicating improper tethering of telomeres to the nuclear envelope and functional loss of CDK2 activity (Fig. 4C). Whereas the lack of OVCAR-3 cytotoxicity from **5** suggests that cyclin-dependent CDK2 activity can be relatively unaffected in somatic cells by these inhibitors, the mouse testicular explant data show the promise of this allosteric CDK2 inhibitor series to disrupt CDK2/SPY1-driven meiosis to yield an efficacious yet safe contraceptive agent.

**Fig. 4.**
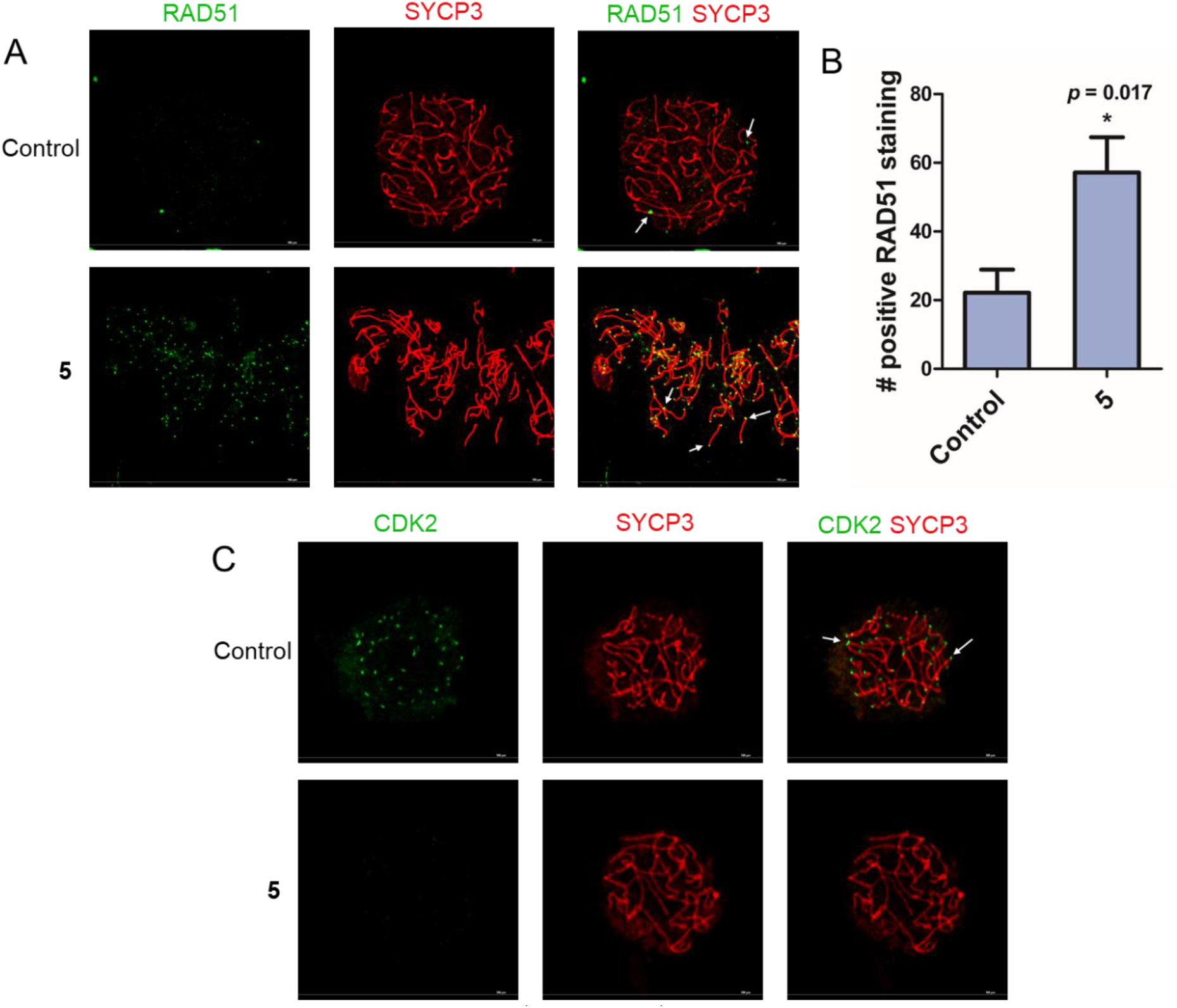
Compound 5 recapitulates features of *Cdk2*^*-/-*^ and *Spdya*^*-/-*^ phenotypes in spermatocytes. A. RAD51 (green) and SYCP3 (red) are stained in explants treated with either control or **5**. B. Compound **5** treated samples show significantly more localization of RAD51 at synapsed regions (positive RAD51 staining at telomeres and chromosomes) compared to control. Values graphed are mean ± SEM (n=6 for each). C. CDK2 (green) and SYCP3 (red) are stained in explants treated with either control or **5**. White arrows added to emphasize the altered localization of RAD51/CDK2 on chromosomes.

## Discussion

The allosteric CDK2 inhibitors in this work stabilize CDK2 in a unique inactive conformation, as demonstrated by X-ray crystallography. Compared to prior structurally-confirmed allosteric CDK2 inhibitors, the affinity of these compounds is markedly improved, demonstrating a K_D_ for CDK2 in the nanomolar range.^17^ This makes these inhibitors the strongest binding, structurally confirmed allosteric CDK inhibitors reported. Furthermore, the disruption of CDK2 PPI by these inhibitors is well characterized by their negative cooperative relationship with cyclin binding and deviates mechanistically in comparison to traditional type I inhibitors like **dinaciclib**. In both biophysical and cellular contexts, these compounds show an impressive selectivity for CDK2 over structurally similar CDK1, a quality not possessed by many previously developed CDK2 inhibitors. The therapeutic potential of these compounds is exemplified by incubation in mouse testicular explants, where they appear to arrest meiosis by recapitulating a phenotype similar to what is observed in *Cdk2*^*-/-*^ and *Spdya*^*-/-*^ spermatocytes.

One of the most striking findings of this series is the marked selectivity for CDK2 over CDK1, a protein with 65% overall sequence identity and 89% sequence similarity in the ATP-site where most CDK2 inhibitors bind.^34^ Historically, the allure of allosteric kinase inhibitors has been their potential for improved selectivity as compared to traditional ATP-site inhibitors due to less structural similarity at pockets distal to the ATP-site. Interestingly, the residues engaged by the anthranilic acid-derived allosteric inhibitors in this paper are nearly identical between CDK1 and CDK2, and are relatively conserved across the CDK family. For example, the carboxylate of the anthranilic acid binds the catalytic lysine (K33) in CDK2, a residue conserved across kinases. An alternative explanation may be the divergent energy landscapes of inactive kinases. Prior work suggests that despite the structural similarity between CDK2 and CDK1, the energy landscapes of these kinases without cyclin bound differ greatly compared to their respective cyclin-bound states.^2^ Because we optimized our ligands to bind an inactive, non-cyclin bound state of CDK2, we observe selectivity in the Jurkat cell CETSA and SPR experiments, where our compounds bind CDK2 but not CDK1. While the ligand-bound inactive states of CDK2 and CDK1 are likely nearly identical due to the structural similarity of these kinases, the binding of these ligands may be biased towards CDK2 over CDK1 due to CDK2’s propensity to sample this inactive state or an enhanced flexibility to accommodate allosteric ligand binding, although the precise mechanism remains to be determined. This latter explanation would be consistent with the large degree of conformational sampling observed for inactive, monomeric CDK2.^26^ Additionally, it has been inferred that inactive CDK1 is more stable than inactive CDK2 and binds cyclin A less readily as a result.^35^

Whereas most previously developed ATP-site inhibitors are often positively cooperative with cyclin and bind the CDK2/cyclin complex, our allosteric inhibitors are negatively cooperative with cyclin.^26^ This is confirmed by the structural FRET assay, where the K_D_ of compound **5** towards CDK2 weakens as more cyclin A is added, and by the kinase activity assay, where the IC_50_ shows less potency as the p-CDK2:cyclin molar ratio decreases. Furthermore, CETSA data with **5** shows stabilization of CDK2 in Jurkat cells, where cyclins are not overexpressed, but not in OVCAR-3 cells, where cyclin E1 drives cell proliferation and survival. While our compounds are not cytotoxic in cyclin overexpressing cancer contexts, disruption of the CDK2/cyclin PPI remains a viable anticancer strategy. Recent work has shown that the complex natural product homoharringtonine (HHT) disrupts the PPI between CDK2 and cyclin A to act as an efficacious anticancer agent, although the allosteric binding mode of HHT has not been experimentally confirmed outside of computational models and its selectivity amongst kinases not explored.^36^ Additionally, disrupting the CDK2/cyclin interaction may be less toxic than traditional ATP-site inhibitors that sequester cyclin via their positive cooperativity with cyclin binding.^16^ Further development of our allosteric inhibitors might confer an advantage in this regard compared to other promising compounds, such as recently developed CDK2/cyclin E1 inhibitor PF-07104091, a selective ATP-site inhibitor targeting the cyclin-bound activated state of the kinase.^37^ Future iterations of our allosteric inhibitors with improved affinity and slower off-rates may very well be selective and efficacious in inhibiting a CDK2/cyclin-driven disease state.

Compound **5** potently affects meiotic CDK2 function, most likely by interfering with SPY1 binding. When the scaffolding and catalytic functions of the CDK2/SPY1 complex are disrupted, prophase I is arrested.^15^ Our inhibitors trigger similar effects, as evidenced by the significantly increased localization of double-strand break repair protein RAD51 on synapses. Additionally, spermatocytes from testis explants treated with **5** show more abnormal CDK2 localization compared to the control. While our compounds arrest meiosis, their potential as effective but safe non-hormonal contraceptive agents requires further exploration. These allosteric inhibitors may selectively disrupt the meiotic CDK2/SPY1 PPI while leaving the mitotic CDK2/cyclin PPI intact. If this selective disruption of CDK2 complexes is true, **5** would be a rare example of a PPI inhibitor that derives its selectivity by exploiting the different affinities of protein binding partners that engage at the same binding site. Aside from the impact of this work on CDK2 inhibitor development, we hope as the conformational landscapes of other kinases are further elucidated and more allosteric pockets are discovered, this work can serve as a model for allosteric inhibitor development to target other therapeutically important kinases.

## Methods

**Dinaciclib** was purchased from Cayman Chemicals with >98% purity. The production and purity assessment (>95%) of GST-CDK1 was made possible via NIH HHSN27520180007I. The data generated for SI Fig. 4 was provided by Reaction Biology via their kinase activity assay and the data generated for Fig. 3B and SI Fig. 10 was provided by Pharmaron via their cytotoxicity assays. The purity of compounds **1**-**5** were determined to be >95% pure by qNMR^38-39^ and described further in the SI.

### CDK2 and Cyclin A Protein Expression and Purification

The gene encoding human CDK2 (1-298) was custom-synthesized (GenScript), subcloned into pGEX6P1 vector providing an *N*-terminal GST-tag and expressed in *E. coli* BL21 DE3 RIPL cells. Cells were grown at 37 °C for 3 h to OD_600_= 0.6, then grown further until OD_600_= 0.9 at 16 °C, induced with 0.5 mM IPTG and allowed to grow further overnight. Cells were harvested by centrifugation at 6000 x *g* for 20 min, the lysate was subjected to GST-affinity chromatography (GE Life Sciences), the GST tag was removed by PreScission protease, and cleaved CDK2 was subjected to a second GST-affinity chromatography step as described previously.^40^ Flow-through fractions containing CDK2 were combined, concentrated and further purified by size exclusion chromatography using a Superdex 75 26/60 column (GE Life Sciences) in 50 mM HEPES, pH 7.5, 150 mM NaCl, 10 mM MgCl_2_, 1 mM EGTA, 0.01 mM ADP, 2 mM DTT. For ITC experiments, purified CDK2 was concentrated to 50-100 μM and flash frozen in appropriate ITC volume aliquots for –80 °C storage.

Expression and purification of p-CDK2 and cyclin A have been described previously.^26^ Briefly, to generate CDK2 phosphorylated on T160 (p-CDK2), a 6His-TEV-CDK2 construct (pCDFduet) was coexpressed with yeast CAK (pGEX) overnight in Terrific broth at 18 °C. Cells were harvested by centrifugation at 5000 x *g* for 20 min and lysed by homogenization (Avestin Emulsiflex C3). Lysates were cleared by centrifugation at 20,000 x *g* for 1 h, subjected to Ni purification (GE HisTrap HP), and p-CDK2 was eluted with 1x PBS, 500 mM imidazole, 10% glycerol. Imidazole was removed by desalting the sample into 1x PBS, 10% glycerol (GE HiPrep 26/10), and the 6His tag was cleaved overnight with TEV protease. The protein was further purified by size exclusion chromatography (GE Superdex S75 10/300 GL), flash frozen and stored at –80 °C.

Bovine cyclin A (171-432 with a *C*-terminal 6His tag) was expressed overnight in BL21(DE3)pLysS in Terrific broth at 20 °C. Cells were lysed and lysates cleared as above, and the protein was isolated through Ni purification into 50 mM Tris pH 8.25, 300 mM NaCl, 100 mM MgCl_2_, 10% glycerol using a 0-500 mM imidazole gradient. Cyclin A was further purified by size exclusion chromatography (GE Superdex 200 16/600) into 50 mM Tris pH 8.25, 100 mM MgCl_2_, 5 mM 2-mercaptoethanol.

### Crystallization

Purified CDK2 was buffer exchanged into 100 mM Na/K phosphate, pH 6.2, 2 mM DTT using PD10 columns (GE Life Sciences) and concentrated to 9 mg/mL. The CDK2/compound **1** complex was obtained by co-crystallization (15% (v/v) Jeffamine ED2001, 50mM HEPES, pH 7.5, 4.5 mg/mL CDK2, 5 mM **1**) at 19 °C. The CDK2/compound **2** complex was obtained by in-diffusion of ligand-free CDK2 crystals with **2** at 19 °C for 24 h (10%(v/v) PEG 3350, 50mM HEPES, pH 7.5, 20% (v/v) ethylene glycol, 4.5 mg/mL CDK2, 5 mM **2**). Both compounds yielded 1 crystal each for diffraction.

### Data Collection and Refinement

X-ray diffraction data of CDK2 liganded with compound **1** was collected on a Rigaku Micro-Max 007-HF X-ray generator at wavelength 1.54178 Å, equipped with a CCD Saturn 944 system located in the Chemical Biology Core of the Moffitt Cancer Center. Data of CDK2 liganded with compound **2** was collected at the GM/CA beamline (23ID-B with Dectris PILATUS 3-6M detector, 23ID-D with Dectris EIGER 16M detector) at Argonne National Laboratory and at wavelength 1.03318 Å. All data were collected at 93.2 K and processed with XDS and Aimless of the CCP4 suite.^41^ Initial phasing of all structures was performed by molecular replacement using PDB 4KD1 as the model in PHASER.^42^ The inhibitor geometrical restraints were obtained with Phenix Elbow.^43^ All structures were refined with Phenix,^44^ and model building was performed with Coot.^45^ Data collection and refinement statistics are shown in Table S1.

### Compound synthesis and characterization

The NMR spectra for the compounds described below are shown in SI Fig. 12-16. The qNMR purity assessment is also included in the SI.

#### General method of S_n_Ar followed by hydrolysis to yield anthranilic acids 1-5

Commercial methyl 2-bromo or 2-fluoro-5-(nitro) or 5-(trifluoromethyl)benzoates (0.30 mmol) were dissolved in DMF (5 mL) with 1.1 equivalents of commercial tryptamine derivatives (in their neutral form or as HCl salts) and sodium bicarbonate (3-4 equiv). An exception was that for **3**, the synthesis of the nitrile tryptamine precursor was accomplished using a previously described method.^46^ The reaction temperature employed was room temperature for compound **1** and 70 °C for compounds **2**-**5**. The solution was stirred for 12 h. After, 1 M HCl (aq) was added until pH < 7 the solution was extracted into DCM (2x). The DCM solvent was removed via rotary evaporation and the crude product was dissolved in MeOH to proceed to the hydrolysis step without purification. Water was added to the MeOH mixture until a 1:1 ratio of solvents was reached. A solution of 5 M NaOH (aq) (1 mL) was added, and the hydrolysis proceeded at 50 °C for 12 h and was monitored by TLC for completion. The mixture was cooled to room temperature and the solvent was removed by rotary evaporation. 1 M HCl (aq) was added until pH < 4 and then the solution was extracted into EtOAc (2x). The EtOAc layer was dried with MgSO_4_ and loaded onto Celite. Subsequent purification was accomplished by silica gel flash column chromatography with a gradient of EtOAc in hexanes of 0-100% over 7 min, eluting around 60% EtOAc as the free carboxylic acid. The solvent was removed by rotary evaporation and the acid was isolated as the final product.

#### 2-((2-(1*H*-Indol-3-yl)ethyl)amino)-5-nitrobenzoic Acid (1)

78% yield. Yellow solid. m.p. 216-218 °C. ^1^H NMR (400 MHz, DMSO-d_6_) δ 13.35 (s, 1H), 10.90 (s, 1H), 8.85 (s, 1H), 8.63 (s, 1H), 8.16 (d, *J* = 9.3 Hz, 1H), 7.60 (d, *J* = 7.8 Hz, 1H), 7.35 (d, *J* = 8.1 Hz, 1H), 7.25 (s, 1H), 7.08 (t, *J* = 7.5 Hz, 1H), 6.99 (t, *J* = 7.4 Hz, 1H), 6.93 (d, *J* = 9.6 Hz, 1H), 3.64 (q, *J* = 6.9 Hz, 2H), 3.07 (t, *J* = 7.0 Hz, 2H).

#### 2-((2-(1*H*-Indol-3-yl)ethyl)amino)-5-(trifluoromethyl)benzoic Acid (2)

33% yield. Light yellow solid. m.p. 201-202 °C. ^1^H NMR (400 MHz, DMSO-d_6_) δ 13.05 (s, 1H), 10.88 (s, 1H), 8.34 (s, 1H), 8.02 (s, 1H), 7.63 (d, *J* = 9.0 Hz, 1H), 7.59 (d, *J* = 7.9 Hz, 1H), 7.35 (d, *J* = 8.1 Hz, 1H), 7.23 (s, 1H), 7.08 (t, *J* = 8.2 Hz, 1H), 7.01 – 6.92 (m, 2H), 3.55 (t, *J* = 7.2 Hz, 2H), 3.05 (t, *J* = 7.0 Hz, 2H).

#### 2-((2-(6-Cyano-1*H*-indol-3-yl)ethyl)amino)-5-(trifluoromethyl)benzoic Acid (3)

9% yield over 5 steps.^46^ Colorless solid. m.p. 190-191 °C. ^1^H NMR (400 MHz, DMSO-d_6_) δ 13.13 – 13.08 (m, 1H), 11.50 (s, 1H), 8.37 (s, 1H), 8.01 (s, 1H), 7.85 (s, 1H), 7.78 (d, *J* = 8.2 Hz, 1H), 7.63 (dd, *J* = 8.9, 2.4 Hz, 1H), 7.57 (s, 1H), 7.32 (dd, *J* = 8.2, 1.5 Hz, 1H), 6.96 (d, *J* = 9.0 Hz, 1H), 3.56 (t, *J* = 7.0 Hz, 2H), 3.07 (t, *J* = 7.0 Hz, 2H).

#### 2-((2-(6-Chloro-1*H*-indol-3-yl)ethyl)amino)-5-(trifluoromethyl)benzoic Acid (4)

70% yield. Colorless solid. m.p. 207-210 °C. ^1^H NMR (400 MHz, DMSO-d_6_) δ 13.09 (s, 1H), 11.03 (s, 1H), 8.33 (s, 1H), 8.01 (s, 1H), 7.67 – 7.57 (m, 2H), 7.39 (s, 1H), 7.29 (s, 1H), 6.99 (dd, *J* = 8.4, 1.9 Hz, 1H), 6.96 (d, *J* = 9.0 Hz, 1H), 3.57 – 3.53 (m, 2H), 3.03 (t, *J* = 7.0 Hz, 2H).

#### 2-((2-(6-Bromo-1*H*-indol-3-yl)ethyl)amino)-5-(trifluoromethyl)benzoic Acid (5)

44% yield. Colorless solid. m.p. 213-214 °C. ^1^H NMR (400 MHz, DMSO-d_6_) δ 13.10 (s, 1H), 11.04 (s, 1H), 8.33 (s, 1H), 8.01 (s, 1H), 7.63 (d, *J* = 8.9 Hz, 1H), 7.60 – 7.51 (m, 1H), 7.28 (s, 1H), 7.11 (d, *J* = 8.4 Hz, 1H), 6.96 (d, *J* = 9.0 Hz, 1H), 3.57 – 3.52 (m, 2H), 3.03 (t, *J* = 7.1 Hz, 2H).

### Isothermal Titration Calorimetry

CDK2 was thawed at room temperature and the buffer was exchanged using Zeba™ spin desalting columns. The final ITC buffer was 1x PBS, 10 mM MgCl_2_, and 5% glycerol (pH = 7.4), with 5% DMSO added after buffer exchange into the protein solution and from a 1:20 concentrated DMSO stock for the ligand solution. CDK2 concentration was determined using a NanoDrop™ spectrophotometer and calculated using absorption at 280 nm (ε = 36,900 M^-1^ cm^-1^). Into a 96-well plate for automatic injection for the MicroCal iTC200 (Malvern), a final volume of 400 μL protein solution for the sample cell and 200 μL inhibitor solution for the injecting syringe were placed into the appropriate wells. The final inhibitor concentration in the syringe was 500-1,000 μM depending on compound solubility. For compounds **3-5**, to get more accurate data on the ITC with appropriate c values,^47^ the CDK2 concentration was lowered ∼10-fold to 5 μM and the syringe concentration lowered to 75 μM. The ITC experiments were conducted at 25 °C and 750 rpm stirring, with a discarded initial 0.4 μL injection and 20 subsequent 4 μL injections with 180 sec between injections. A one-site binding model was used to fit the ITC data after adjusting the baseline to account for slight buffer mismatch between the cell and syringe samples.

### Surface Plasmon Resonance (SPR)

The SPR method used was largely adapted from a published method for CDK1 and CDK2 SPR studies.^2^ A Biacore S200 (Cytiva) at 20 °C and CM5 chips were used for binding analysis. Multi-cycle runs were used for small molecule binding analysis, whereas a single-cycle run was used to measure cyclin binding. The buffer used to prepare the protein samples was 20 mM HEPES, 150 mM NaCl, 10 mM MgCl_2_, 0.01% Tween 20, pH = 7.4). For runs with small molecules, an additional 1% DMSO was added for solubility. GST capture kit conditions (Cytiva catalog number BR100223) were used to capture anti-GST antibody on both the sample and reference cells (7 min immobilization, 10 μL/min flow rate). Both surfaces had high affinity sites capped with an additional 3 min of GST flowed over (5 μg/mL concentration, 5 μL/min flow rate) followed by regeneration (10 mM glycine, pH = 2.2). On the subtractive reference surface, GST was immobilized (20 μg/mL, 5 μL/min, 5 min). On the sample surface, GST-tagged CDK1 or GST-tagged CDK2 (with the GST cleavage stepped skipped during the purification process) was immobilized in a similar fashion (50 μg/mL, 5 μL/min, 5 min). For GST-tagged unphosphorylated CDK1, a filtration through a Fisherbrand™ syringe filter: PTFE Membrane (0.45 *μ*m pore size, 13 mm diameter) was performed to lessen aggregation.

For small molecule binding analysis, two startup cycles of running buffer were carried out on both surfaces after protein immobilization. Depending on observed binding kinetics, a 60-180 sec association time, and a 180-360 sec dissociation time at 30 μL/min for each concentration of a particular compound over both surfaces was injected. An eight-point dose response was conducted with increasing concentrations for each compound with concentrations that were >10*K_D_ at the highest and <0.1*K_D_ at the lowest. To evaluate the binding of **5** against both CDK1 and CDK2, the concentrations employed were based on its K_D_ for CDK2, as **5** did not appear to bind CDK1. For **1**, the highest concentration used was 25 μM due to nonspecific binding to the chip surface at higher concentrations. Zero concentrations were performed at the beginning and end of the dose response as a baseline and to ensure internal reproducibility. An additional maximum concentration of compound was injected after the second zero concentration as an additional internal control. Lastly, a five-point DMSO solvent correction was performed for each compound before analysis with 0.6%, 0.8%, 1.0%, 1.2%, and 1.4% DMSO in buffer. To determine the K_D_ and kinetic binding parameters of each compound against their respective protein, the Biacore S200 Evaluation Software (Cytiva) using affinity fit via the steady-state approximation (SSA), and when possible kinetic fit (k_off_/k_on_), was used.

For protein-protein affinity analysis with cyclin, the same buffer was used but without DMSO. A single-cycle run with increasing concentrations of cyclin A was used to determine the affinity of the CDK2/cyclin A interaction due to the slow off-rate of cyclin binding. After protein immobilization, one startup run of buffer was flowed over both the reference and sample surface to equilibrate the system. In an eight-point dose response, each concentration of cyclin (0-100 nM) had a 180 sec association time and 180 sec dissociation time. A kinetic fit was used to determine the K_D_. For both small molecule and cyclin affinity experiments, duplicates were conducted and the K_D_ was expressed as the mean ± sample standard deviation.

### Förster resonant energy transfer (FRET) assay

The CDK2 FRET assay between cyclin and small molecules has been described previously.^26^ Briefly, CDK2 (C118A C177S A93C R157C) was labeled with FRET donor and acceptor dyes (AF488, AF568; Fluoroprobes), purified by gel filtration, mixed with varying concentrations of bovine cyclin A and dispensed into a 384-well plate (Greiner Bio-One) into which 1 μL inhibitor had been pre-dispensed. The final concentration of CDK2 in the FRET assays was 5 nM in 1x PBS pH 7.4, 10 mM MgCl_2_, 5 mM DTT, 0.5 mg/mL bovine gamma globulins and 0.02 % Tween 20. Plates were incubated for 30 min, and measurements were performed on a fluorescence plate reader (Fluorescence Innovations). Emission peaks were spectrally unmixed,^48^ and the ratios of the donor and acceptor peaks were used to calculate the apparent K_D_ for each inhibitor.

### ADP-Glo experiments

Kinase activity of phosphorylated CDK2 with cyclin A and varying amounts of inhibitor (**dinaciclib** or **5**) were measured using the ADP-Glo Kinase Assay (Promega) in a luminescence plate reader (Tecan Infinite M1000 PRO). Triplicate reactions were carried out in ADP Quest Assay buffer (15 mM HEPES, pH 7.4, 150 mM NaCl, 1 mM EGTA, 0.02% Tween-20, 10 mM MgCl_2_, and 0.1 mg/mL bovine-γ-globulins). Assays were performed with 5 nM phosphorylated CDK2, 200 μM ATP (Promega), 100 μM PKTPPKAKKL substrate (GenScript), and varying concentrations of cyclin A (5, 50, or 500 nM). Phosphorylated CDK2 was pre-incubated with inhibitor at 20 °C for 15 min. Reactions were initiated by simultaneously adding ATP and cyclin A and incubated at 20 °C for 15 min. Reactions were then quenched with ADP-Glo reagent following the manufacturer’s instruction. Luminescence was measured using a 1 sec integration time and normalized to the no-inhibitor 5% DMSO controls. Kinase activity was determined by globally fitting relative luminescence data from all three cyclin A concentrations to a quadratic binding model using non-linear regression in GraphPad Prism 9. For fitting, CDK2 concentration was constrained to 5 nM with data sets from each p-CDK2:cyclin A ratio globally constrained to share the lower baseline parameter. IC_50_ values reported represent the 95 % confidence intervals obtained from the fits (n=3) using the symmetrical likelihood method in GraphPad Prism 9.

### Cellular Thermal Shift Assay (CETSA)

Jurkat and OVCAR-3 cell lines were purchased from the American Type Culture Collection. Both Jurkat and OVCAR-3 cells were cultured in RPMI 1640 (Corning) medium at 37 °C in a humidified chamber under 5% CO_2_. Medium for Jurkat was supplemented with 10% fetal bovine serum (Gibco) and 1% penicillin-streptomycin (Corning), and OVCAR-3 with 20% fetal bovine serum and 1% penicillin-streptomycin. The CETSA was performed following a reported protocol.^49^ Briefly, in two separate T75 flasks for each cell line, Jurkat cells were grown to ∼3 million/mL in 10 mL medium and OVCAR-3 cells to ∼1 million/mL in 10 mL medium. Compound **5** was added to one flask for each cell line with a final concentration of 20 μM and DMSO (0.5%) was added to the other respective flasks. Cells were incubated with the indicated treatments for 1 h (Jurkat) or 2 h (OVCAR-3) at 37 °C, then collected through centrifugation and resuspended in 1 mL PBS supplemented with complete protease inhibitor cocktail (Roche) and PhosSTOP phosphatase inhibitor cocktail (Roche). For each treatment group, cells were aliquoted into ten different PCR tubes with 100 μL of cell suspension in each tube (∼3 million cells per tube for Jurkat and ∼1 million cells per tube for OVCAR-3). Cells in each tube were heated to their designated temperatures (40–67 °C) for 3 min in a thermal cycler, and then the tubes were incubated at room temperature for another 3 min before snap-freeze of the samples in liquid nitrogen. Cells were fully lysed by two more cycles of freeze-thaw. The lysates were cleared by centrifugation, and an equivalent amount (12 μL) of cell lysate from each sample was electrophoresed on a 4-12% NuPAGE gradient gel (Invitrogen) and transferred onto low florescent polyvinylidene difluoride membranes (Bio-Rad). Immunoblotting was performed with anti-CDK2 (sc-6248, Santa Cruz Biotechnology, 1:2,000), anti-CDK1 (sc-54, Santa Cruz Biotechnology, 1:2,000) and anti-β-actin (A1978, Millipore Sigma, 1:2,000) antibodies followed by secondary goat-anti-mouse (A16072, Invitrogen, 1:1,000 for CDK2 and CDK1 blots) and goat-anti-mouse (A32729, Invitrogen, 1:1,000 for β-actin blots) Alexa680 antibodies. When using horseradish peroxidase-conjugated anti-mouse antibody for CDK2 and CDK1, West Femto Maximum Sensitivity Substrate (Thermo Scientific) was added to the membranes before imaging on an Odyssey Fc Imaging system (Li-Cor). Band intensity was quantified with ImageJ 1.52a, and data were then fitted to obtain apparent T_agg_ values using the Boltzmann Sigmoid equation within GraphPad Prism (v. 9.1.1) software. Densitometry data in the melting curves are mean ± SEM from three biological replicates.

### Chromosome Spread Preparation

For spermatocyte chromosome spreads, adult WT C57BL/6J mouse testes were dissected and incubated at 37 °C for 6 h in either Dulbecco’s Modified Eagle’s Medium/Nutrient Mixture F-12 (DMEM/F12, Thermo Scientific) or compound **5** (100 μM) in DMEM/F12. Following incubation, meiotic chromosome spreads were prepared as described previously.^50^ Briefly, samples were placed in a hypotonic extraction buffer (30 mM Trizma hydrochloride, 50 mM sucrose, 17 mM sodium citrate, 5 mM EDTA, 0.5 mM DTT, and protease and phosphatase inhibitor cocktail [1:100]; Sigma-Aldrich) for 10 min. Samples were then moved to a 100 mM sucrose solution (pH 8.2) and spermatocytes were detached by pipetting. Drops of the cell suspension were then placed onto a coverslip that had previously been soaked in a 1% paraformaldehyde (Sigma-Aldrich) and 0.15% Triton X-100 (Sigma-Aldrich) solution. Coverslips were dried at room temperature overnight and either used immediately for staining or stored at – 80 °C.

### Immunofluorescent Staining and Analysis

For immunofluorescent staining of chromosome spreads, slides were washed with PBS (Thermo Scientific) and 0.015% Triton X-100, blocked with 5% bovine serum albumin (Sigma-Aldrich) in PBS at room temperature for 1 h, and then incubated with anti-CDK2 monoclonal antibody (1:100) and anti-SCP1 polyclonal antibody (NB300-229, Novus, 1:100) or anti-RAD51 polyclonal antibody (pc130, Millipore, 1:100) and anti-SYCP-3 monoclonal antibody (sc-74569, Santa Cruz Biotechnology, 1:100) overnight at 4 °C. Fluorescence was developed using Alexa Fluor 488 and 564 conjugated secondary antibodies (ab150073, Life Technologies, 1:250). A Nikon Eclipse TiE inverted microscope with A1R-SHR confocal was used for imaging. RAD51 localization was quantitively assessed by determining the number of positive staining sites on the chromosome and telomeres per cell for 10 samples per group. An unpaired two-tail t-test with *p* <0.05 was used to determine significance. CDK2 staining was qualitatively assessed through abnormal or normal scoring for 10 samples per group, where an abnormal score was given if 3+ telomeres within the sample did not stain positive for CDK2.

## Acknowledgements

E.B.F. was supported by NIH/NIGMS (through training grants T32 GM008244 and T32 GM132029), as well as by an NIH/NCI fellowship (F30 CA232303). Funding for this project was provided by NICHD: R01 HD080431 (V.C.), R61 HD099743 (G.I.G. and V.C.), NIGMS: R01 GM121515 (N.M.L.), DOD: W81XWH-21-1-0674 (D.A.H.), and W81XWH-19-1-0336 (J.T.). Production of CDK1 was made possible by NIH HHSN27520180007I. We acknowledge support by the GM/CA beamlines at the Advanced Photon Source (APS), funded by the National Cancer Institute (ACB-12002) and the National Institute of General Medical Sciences (AGM-12006, P30 GM138396, S10 OD012289). The APS is a U.S. Department of Energy Office of Science User Facility operated under Contract No. DE-AC02-06CH11357. The Moffitt Chemical Biology Core is acknowledged for use of the crystallization and X-ray facilities (National Cancer Institute grant P30 CA076292). We thank Dr. Tim Ward for his initial contributions to this project. Reaction Biology provided the data for the CDK activity data in SI Fig. 4 and Pharmaron provided data for SI Fig. 10.

## Author Contributions

E.B.F, J.T., E.R., S.G., L.S., N.W., D.R., A.M., and K.J. designed experiments, acquired data, and analyzed data for this work. A.N. and H.K. acquired data presented in this work. J.E.H., N.M.L., E.S., V.C., D.A.H., and G.I.G. helped conceive and design experiments for this work as well as analyze data.

## Supplementary Information (SI)

Supplementary Information is available for this paper.

## Materials and Correspondence

Correspondence and requests for materials should be sent to Gunda I. Georg at.

