## Supplementary information for "Development of allosteric, selective cyclin-dependent kinase 2 (CDK2) inhibitors that are negatively cooperative with cyclin binding and show potential as contraceptive agents"

SI Fig. 1. Compound **1** binds to CDK2 with a $K_D$ = 1.8 ± 1.2 μM with fast association, fast dissociation kinetics as determined by SPR……………………………………………………S2

SI Fig. 2. ITC traces of **2**, **3, 4** into CDK2……………………………….………………….S2

SI Fig. 3. SPR traces of **4** (A), **5** (B), and **3** (C) into CDK2……………………………………S3

SI Fig. 4. Compound **4** does not substantially inhibit any CDK/cyclin complex at a concentration well above its $K_D$. Data generated by Reaction Biology……………………..………………S3

SI Fig. 5. Compound **4** demonstrates a negatively cooperative relationship with cyclin binding in CDK2……………………………………………………………………………………….S4

SI Fig. 6. **Dinaciclib** inhibits CDK2 independent of cyclin concentration………………………S4

SI Fig. 7. Replicate of SPR experiment measuring cyclin binding to CDK2………………..…..S5

SI Fig. 8. Triplicate CETSA data for DMSO (control) and compound **5** treated Jurkat cells……S6

SI Fig. 9. In an SPR assay, compound **5** does not bind CDK1 (A) whereas **dinaciclib** binds CDK1 with an affinity similar to its published value (B)…………………………………………...S7

SI Fig. 10. Compound **5** shows minimal cytotoxicity against a cell line with cyclin E1 overexpression and dependent on CDK2 activity (OVCAR-3), in contrast to the toxic kinase inhibitor staurosporine with $IC_{50}$ values in the low nanomolar range…….……..……………S7

SI Fig. 11. Triplicate CETSA data for DMSO (control) and compound **5** treated OVCAR-3 cells………………………………………………………………………………………S8

SI Fig. 12. $^1$H NMR spectrum of **1**……………………………………..…………………..…..S9

SI Fig. 13. $^1$H NMR spectrum of **2**……………………………………..……………….….S10

SI Fig. 14. $^1$H NMR spectrum of **3**……………………………………..………………..…S11

SI Fig. 15. $^1$H NMR spectrum of **4**……………………………………..…………….…….S12

SI Fig. 16. $^1$H NMR spectrum of **5**…………………………………………………..…………S13

Table S1. Crystallographic data and refinement statistics…………………..…………………S14

Absolute qNMR data………………………………………………………………….……...S15-S17

Uncropped immunoblotting images ……………………………………………………S18-S23

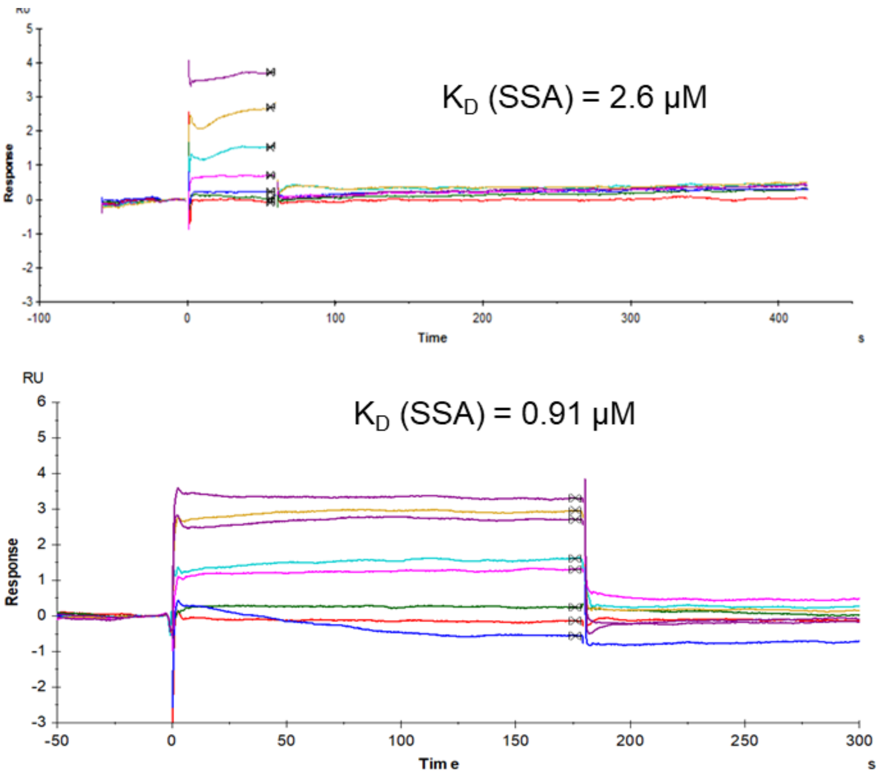

SI Fig. 1. Compound **1** binds to CDK2 with a $K_D$ = 1.8 ± 1.2 μM (n = 2) with fast association, fast dissociation kinetics as determined by SPR.

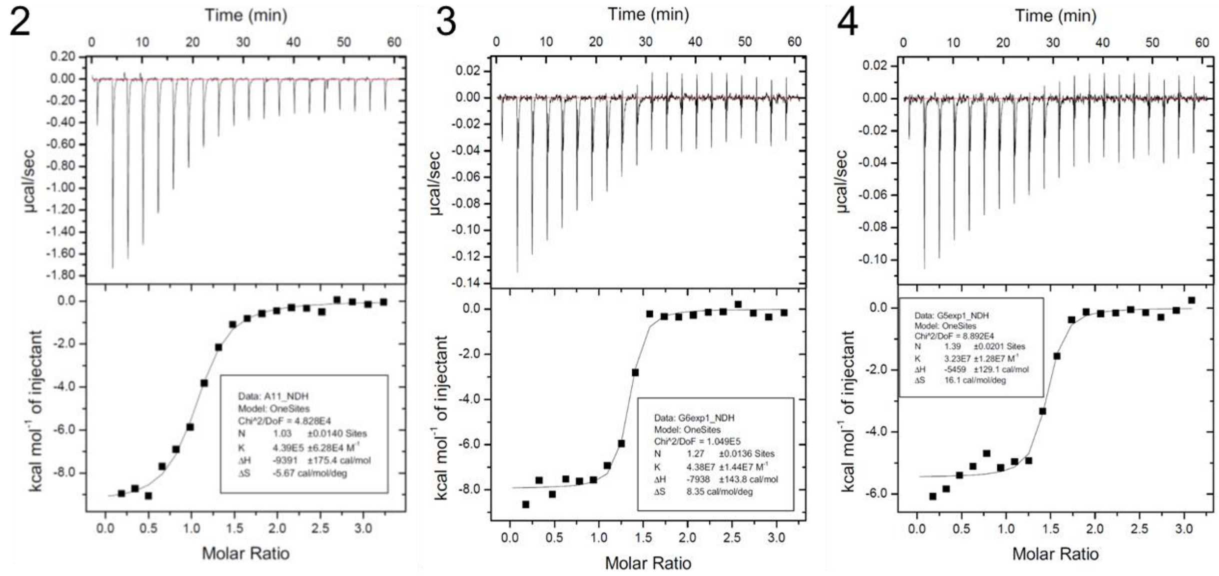

SI Fig. 2. ITC traces of **2**, **3**, and **4** against CDK2. Results summarized in Table 1.

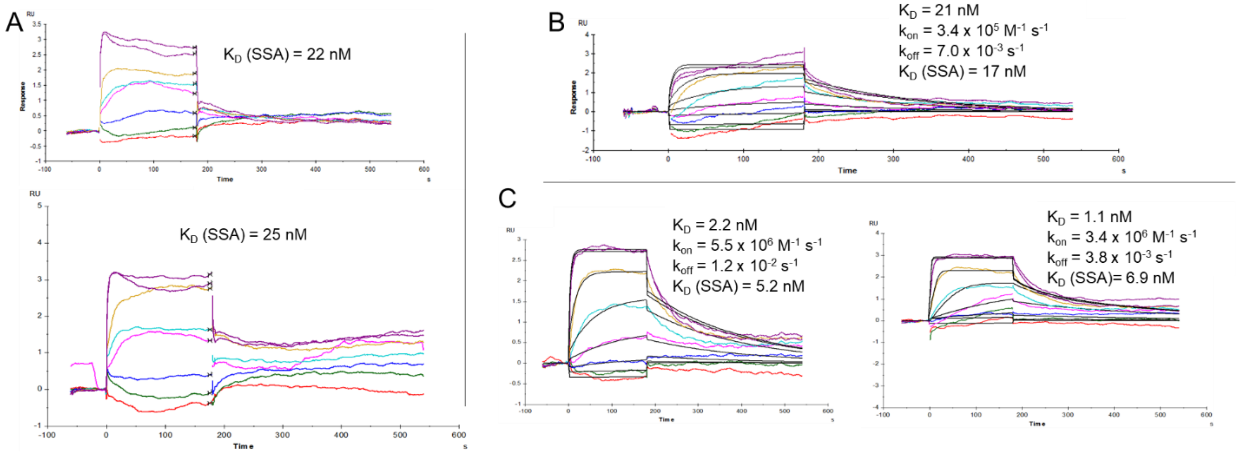

SI Fig. 3. SPR traces of **4** (A, $K_D = 24 \pm 2$ nM, n = 2), **5** (B, $K_D$ (SSA) = $16 \pm 2$ nM, $K_D$ ($k_{off}/k_{on}$) = $19 \pm 2$ nM, n = 2), and **3** (C, $K_D$ (SSA) = $6.1 \pm 1.2$ nM, $K_D$ ($k_{off}/k_{on}$) = $1.7 \pm 0.8$ nM, n = 2) into CDK2. A replicate for **5** is shown in Fig. 2E in the manuscript.

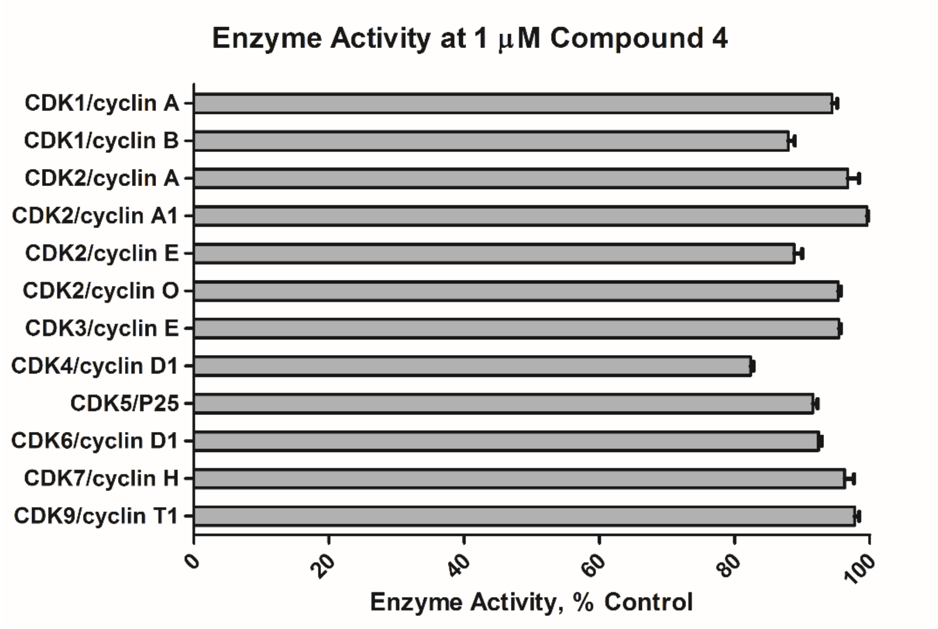

SI Fig. 4. Compound **4** does not substantially inhibit any CDK/cyclin complex at a concentration well above its $K_D$. Data generated by Reaction Biology.

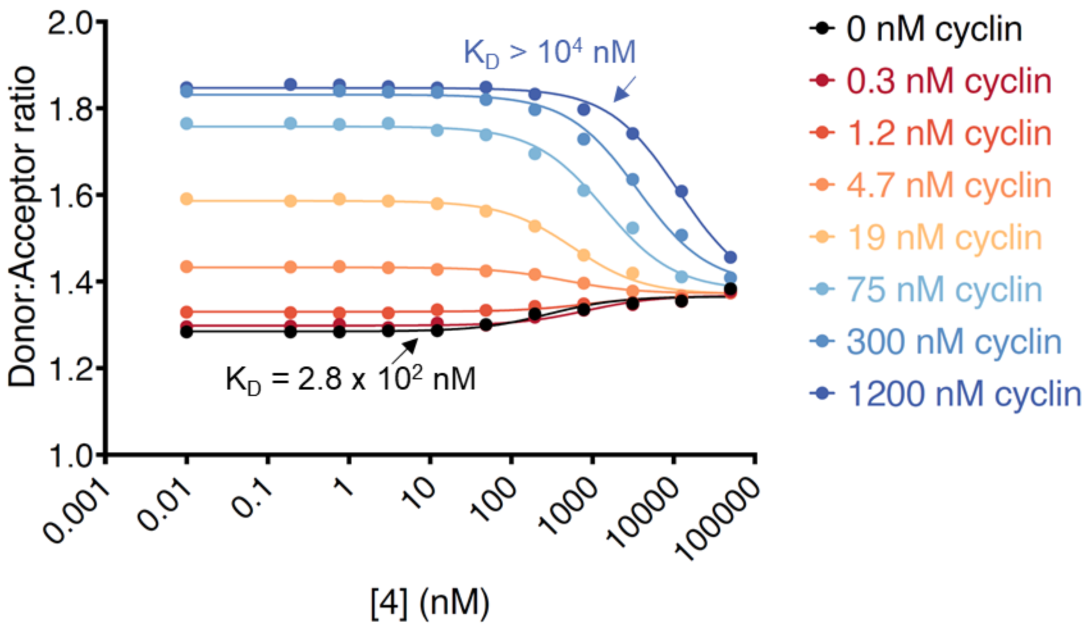

SI Fig. 5. Compound **4** demonstrates a negatively cooperative relationship with cyclin binding in CDK2.

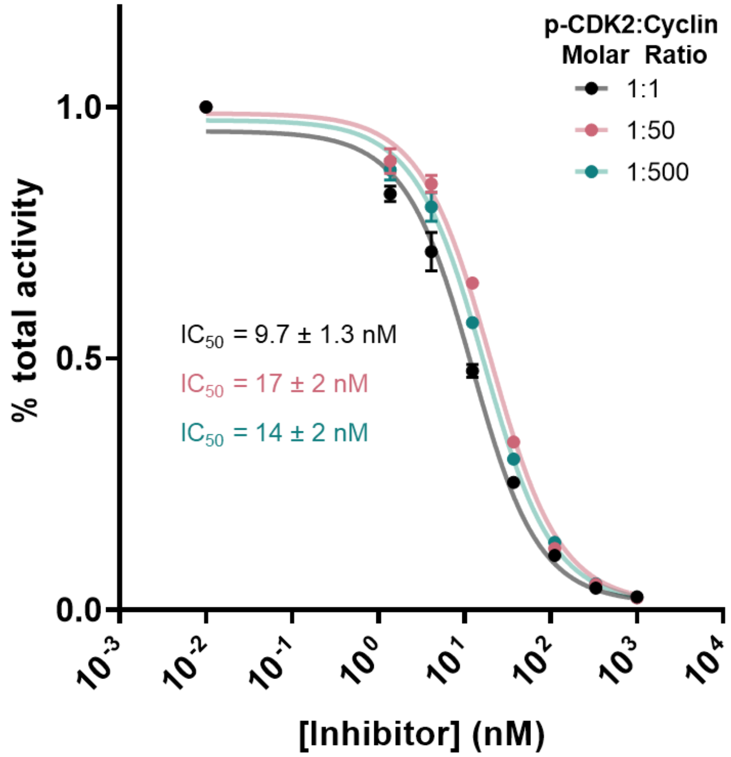

SI Fig. 6. **Dinaciclib** inhibits CDK2 largely independent of cyclin concentration.

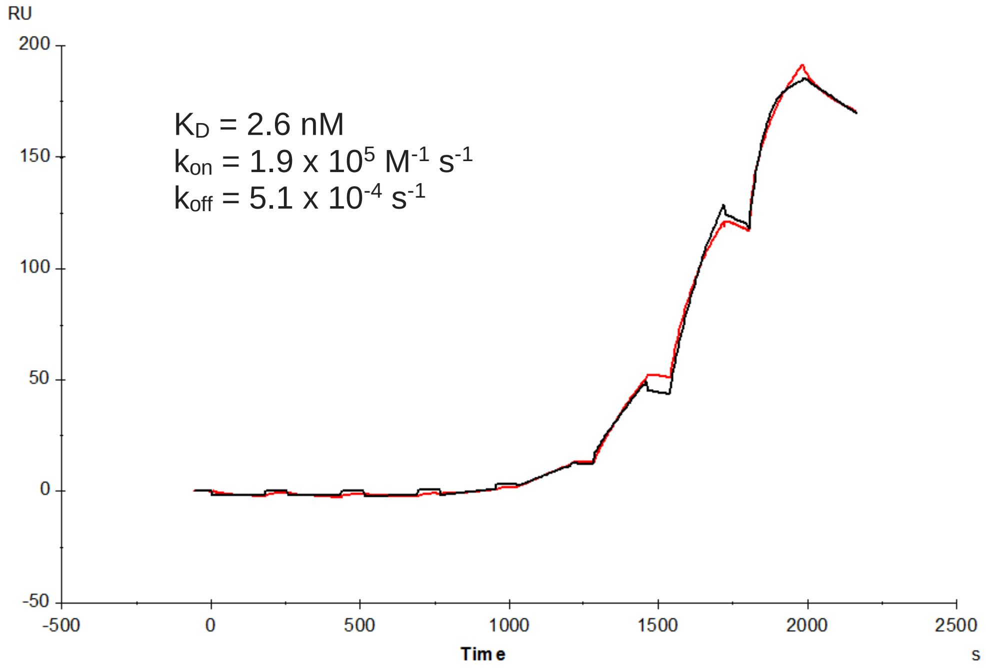

SI Fig. 7. Replicate of SPR experiment measuring cyclin binding to CDK2.

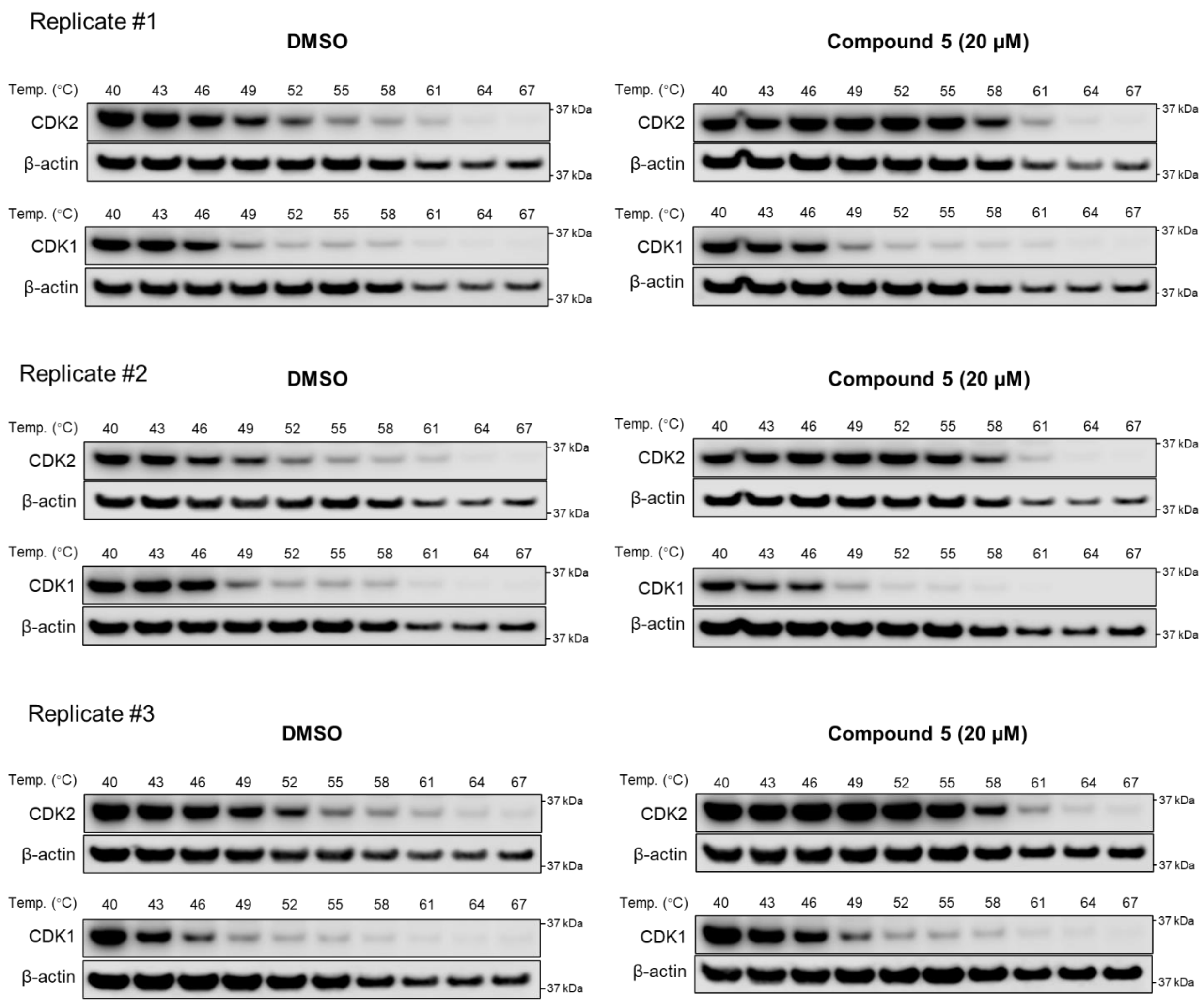

SI Fig. 8. Triplicate CETSA data for DMSO (control) and compound **5** treated Jurkat cells. Numbers shown above bands on blots indicate the temperature of the experiment (°C).

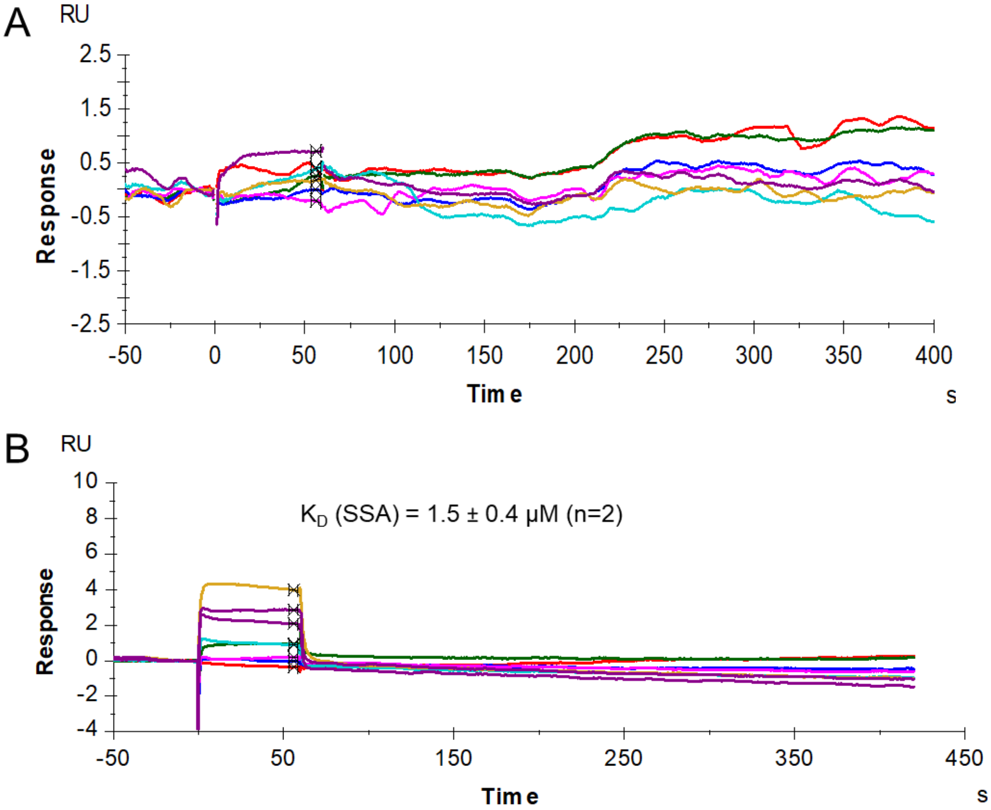

SI Fig. 9. In an SPR assay, compound **5** does not bind CDK1 (A) whereas **dinaciclib** binds CDK1 with an affinity similar to its published value (B, published value[1]: $K_D$ (SSA) = 1.8 ± 0.2 μM). **Dinaciclib** $K_D$ from duplicate runs expressed as mean ± standard deviation.

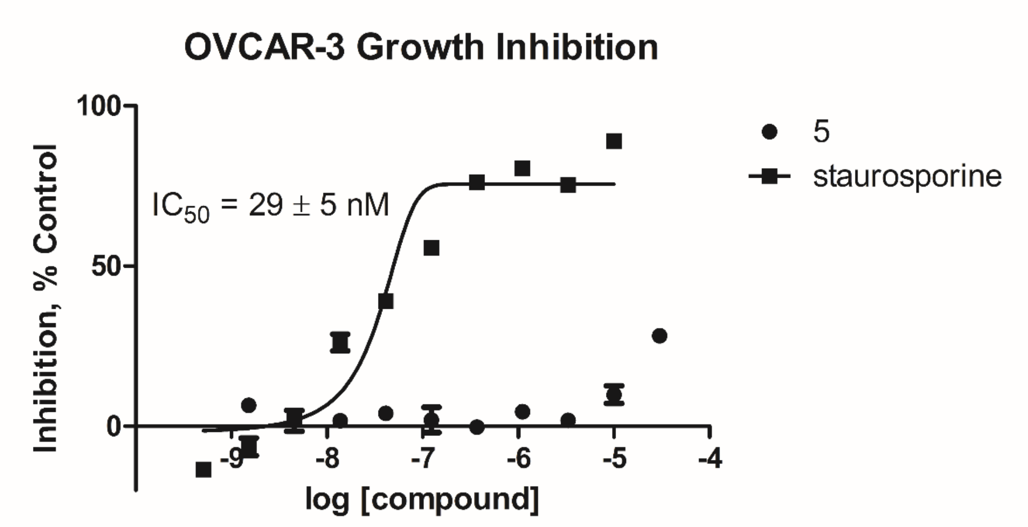

SI Fig. 10. Compound **5** shows minimal cytotoxicity against a cell line with cyclin E1 overexpression and dependent on CDK2 activity (OVCAR-3), in contrast to the toxic kinase inhibitor staurosporine with $IC_{50}$ values in the low nanomolar range.

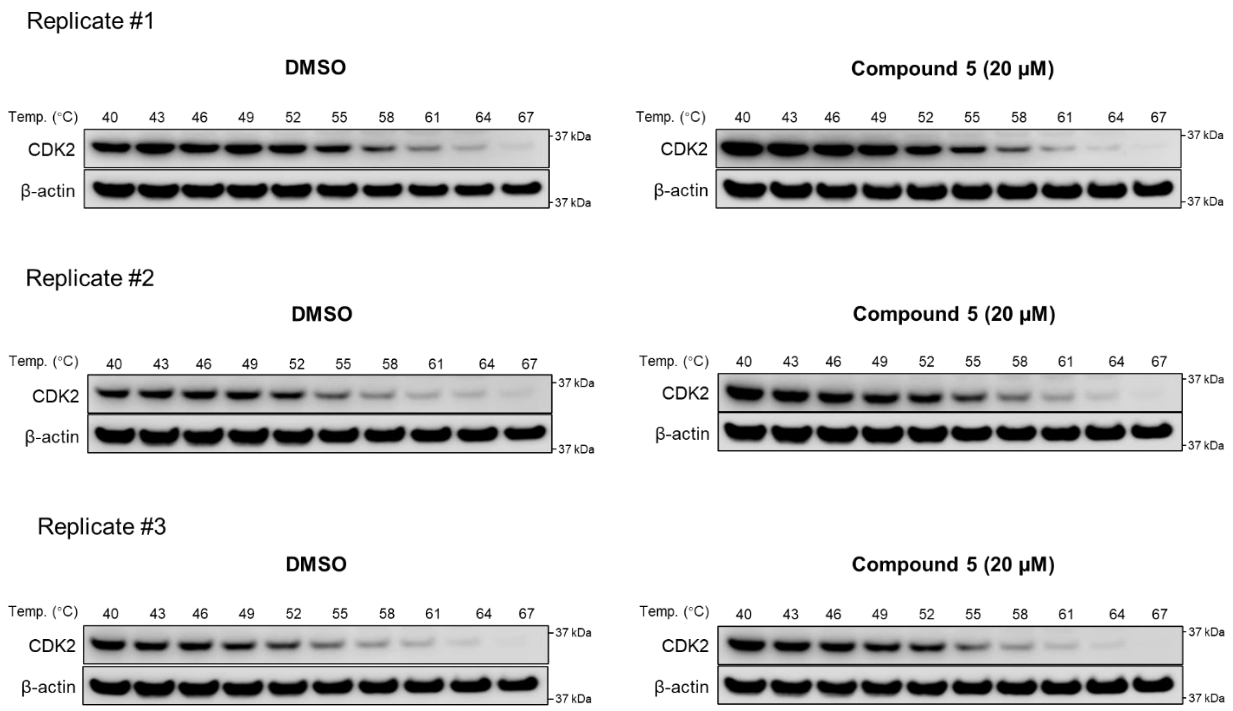

SI Fig 11. Triplicate CETSA data for DMSO (control) and compound **5** treated OVCAR-3 cells. Numbers shown above bands on blots indicate the temperature of the experiment (°C).

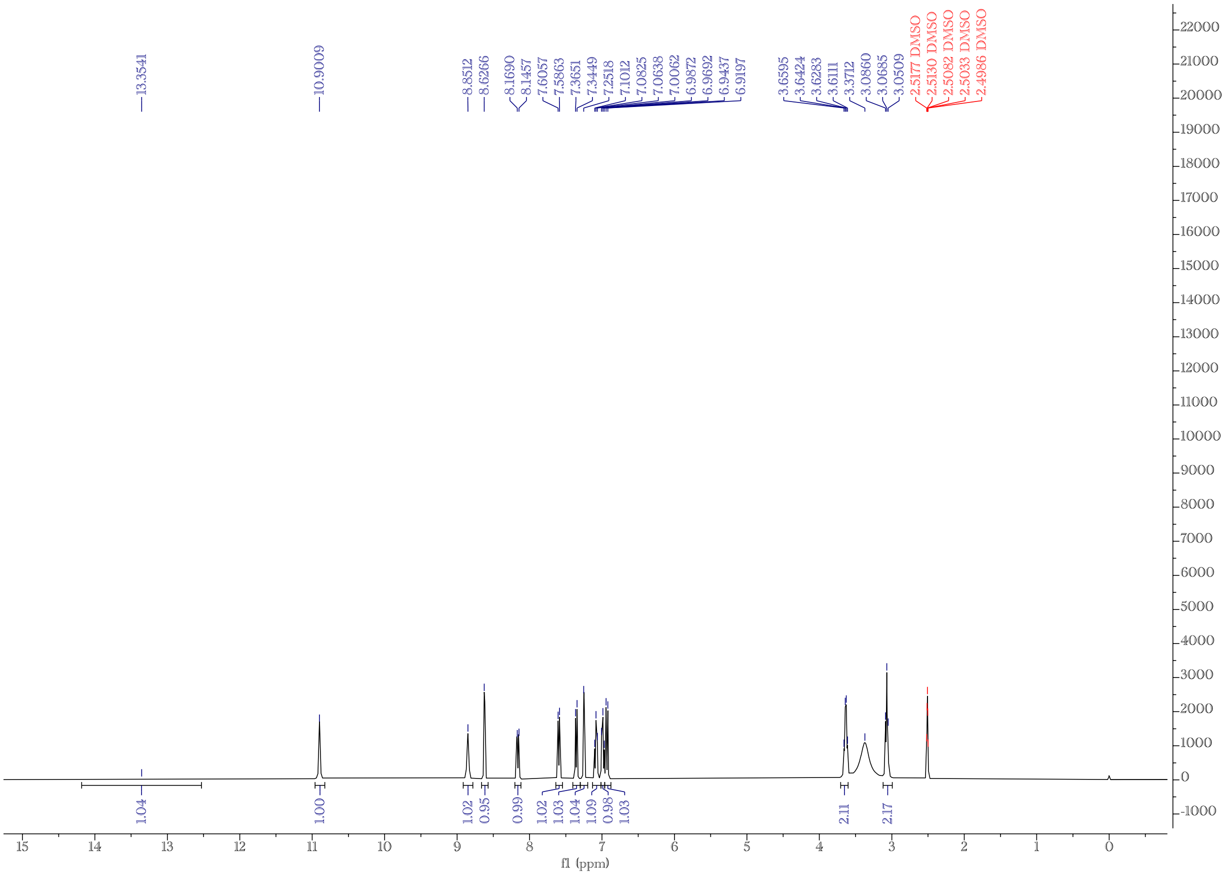

SI Fig. 12. $^{1}$H NMR spectrum of **1**.

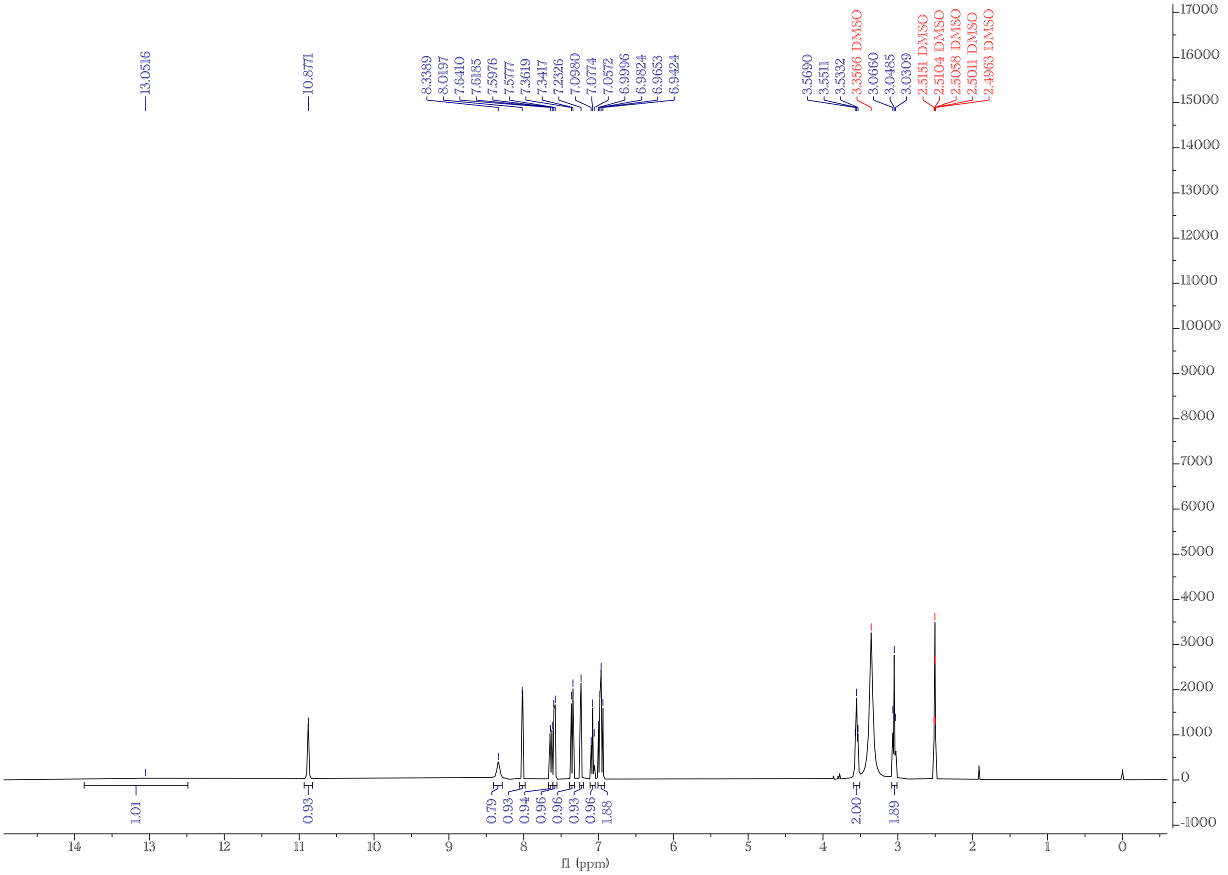

SI Fig. 13. $^{1}$H NMR spectrum of **2**.

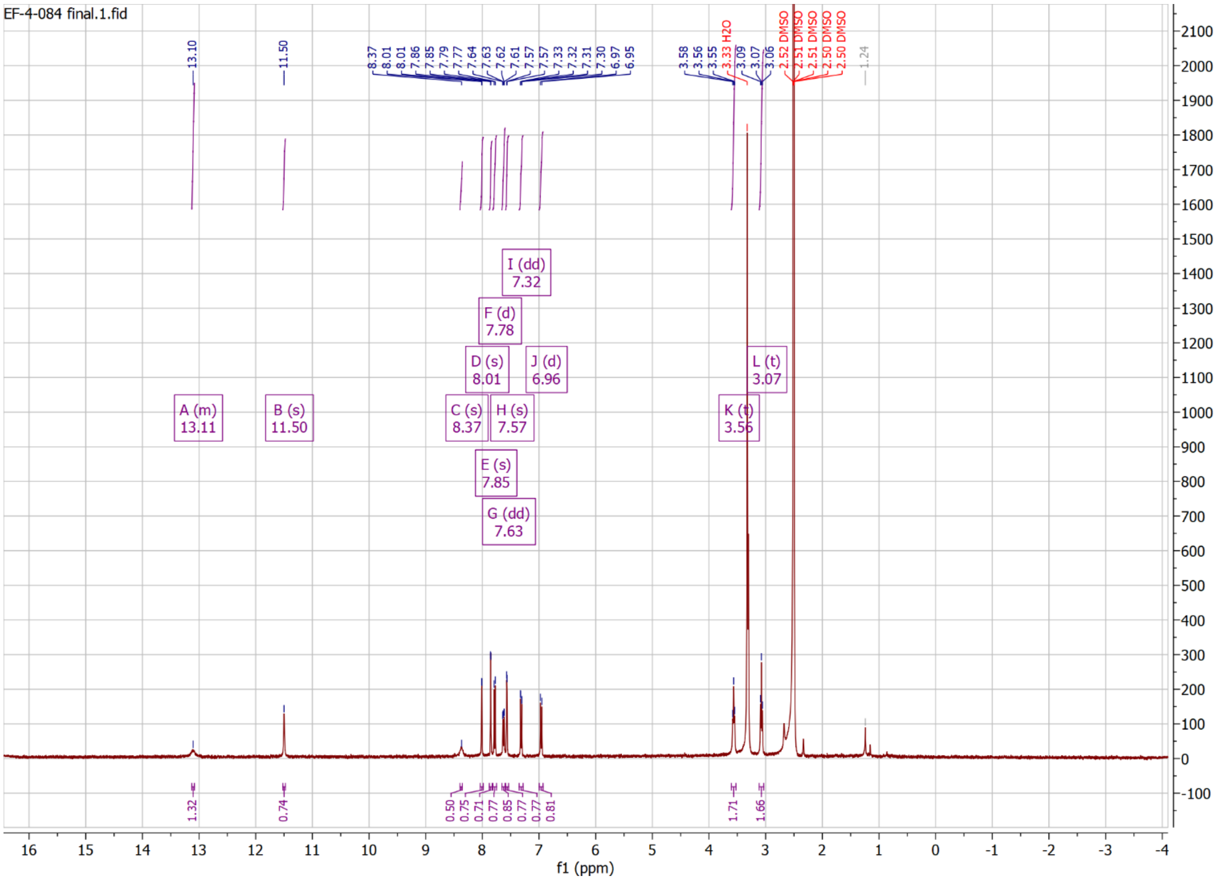

SI Fig. 14. $^{1}$H NMR spectrum of **3**.

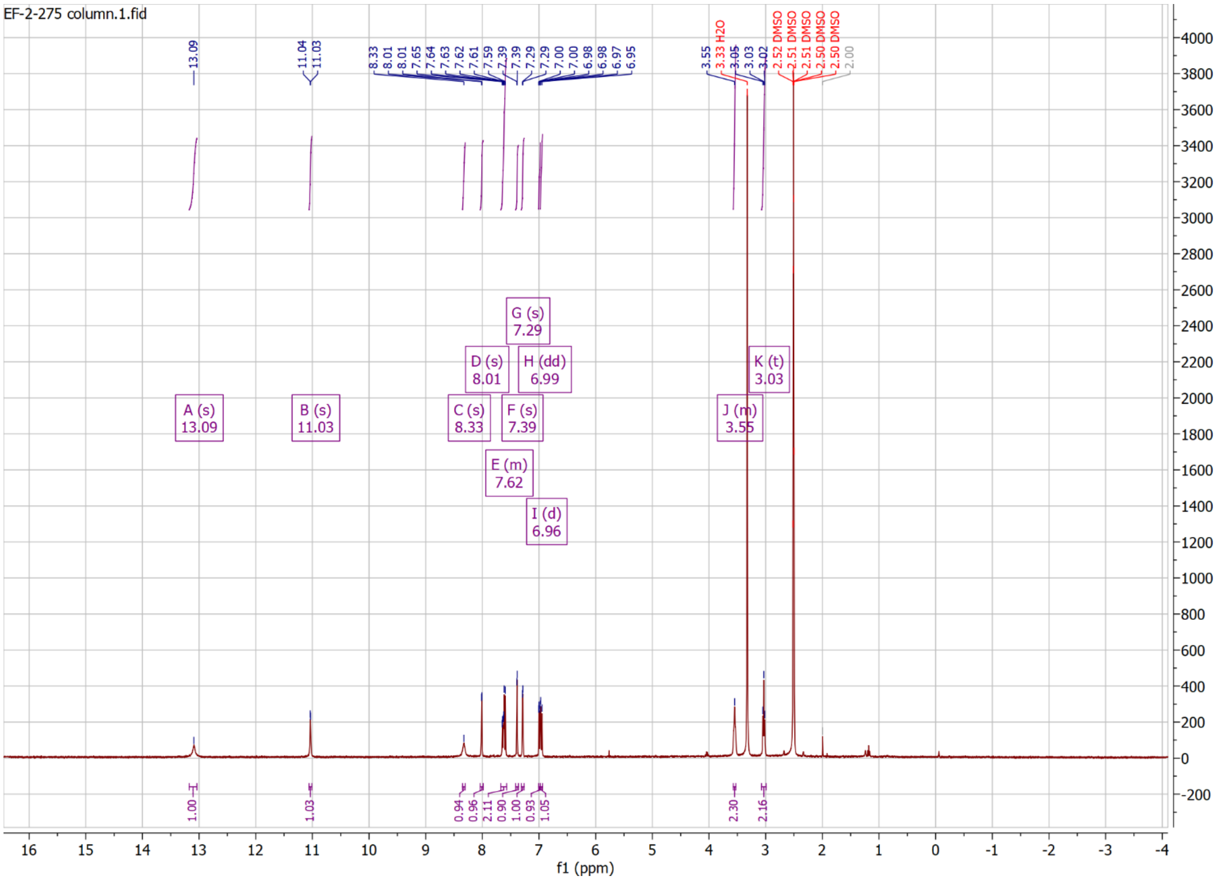

SI Fig. 15. $^{1}$H NMR spectrum of **4**.

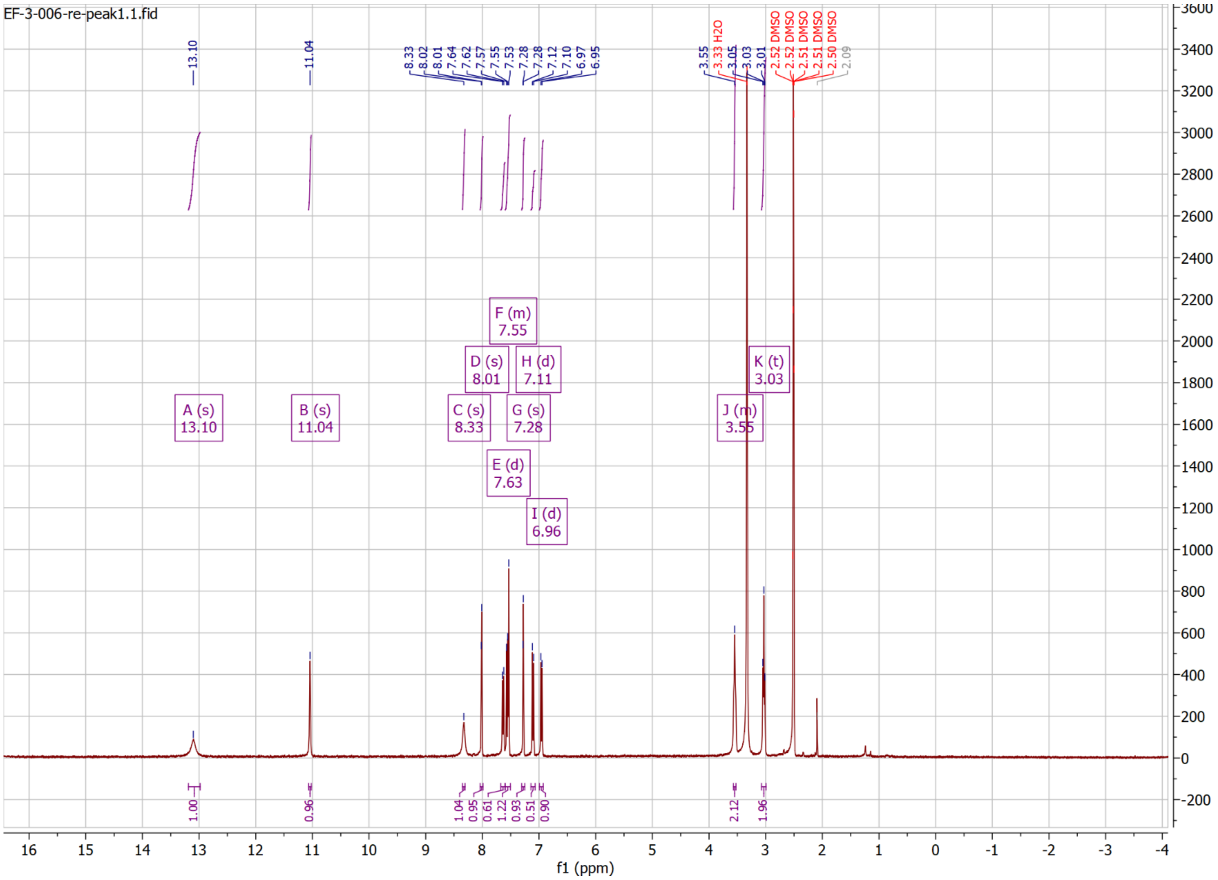

SI Fig. 16. [1]H NMR spectrum of **5**.

**Table S1: Crystallographic data and refinement statistics**

| **Protein** | | | |
|---|---|---|---|
| **Inhibitor** | | Compound **1** | Compound **2** |
| **PDB ID** | | 7RWF | 7S84 |
| **Ligand code** | | 7TW | 8IL |
| **Space group** | | P 21 21 21 | P 21 21 21 |
| **Unit cell dimensions** | a | 50.80 | 54.19 |
| | b | 70.24 | 71.55 |
| | c | 71.44 | 72.22 |
| | α | 90 | 90 |
| | β | 90 | 90.00 |
| | γ | 90 | 90 |
| **Resolution range (Å)** | | 41.4 - 1.5 (1.554 - 1.5) | 43.2 - 2.0 (2.072 - 2.0) |
| **Unique reflections** | | 42774 (4172) | 19444 (1879) |
| **Rmeas (%)** | | 11.3 (47.8) | 11.9 (75.1) |
| **Completeness (%)** | | 99.93 (99.33) | 99.37 (98.74) |
| **I/σI** | | 20.85 (2.02) | 9.77 (2.07) |
| **Rwork (%)** | | 19.22 (20.53) | 19.69 (22.46) |
| **Rfree[a] (%)** | | 20.94 (21.81) | 24.45 (30.87) |
| **Wilson B ($Å^2$)** | | 17.01 | 35.32 |
| **Average B ($Å^2$)** | all | 21.10 | 35.13 |
| | protein | 20.78 | 31.38 |
| | ligand | 19.32 | 33.96 |
| | solvent | 26.09 | 38.81 |
| **rmsd[b] bond lengths (Å)** | | 0.007 | 0.009 |
| **rmsd angles (deg)** | | 0.93 | 1.04 |
| **Ramachandran** | favored (%) | 98.32 | 98.16 |
| | allowed (%) | 1.68 | 1.84 |
| | outliers (%) | 0 | 0 |

Values in paranthesis are for the highest resolution bins.

[a] Rfree is Rcryst calculated for randomly chosen unique reflections, which were excluded from the refinement.

[b] rmsd = root-mean-square deviation from ideal values.

Absolute qNMR data

(1) Sample preparation

Compounds and dimethyl sulfone (*TCI*, 100% by GC, lot VOA5G-OD) were weighed into 3 mm standard NMR tubes using an analytical balance (Mettler Toledo XS205) with 0.01 mg accuracy, then 170 uL of Acetone-$d_6$ was added into NMR tubes. All NMR tubes were capped and wrapped with PTFE tape and subsequently with paraffin tape. The compounds and the internal calibrant were dissolved completely before submitted to NMR instrument.

| Compounds ID | Weight of compounds (mg) | Weight of $DMSO_2$ (mg) |
|---|---|---|
| **1** | 5.70 | 1.65 |
| **2** | 2.44 | 4.01 |
| **3** | 4.07 | 2.62 |
| **4** | 2.12 | 1.91 |
| **5** | 2.81 | 2.28 |

(2) Acquisition
The instrument and controlled parameters used during the acquisition are listed below:

*Pulse Program*: Single pulse, without carbon decoupling (‘s2pul’ [Agilent/Varian]; ‘zg’ with $90^{o}$ pulse [Bruker]; “single pulse” [Jeol])
*Sample Temperature*: 25 °C
*Data Points*: 64 K
*Zero-Filling*: to 256 K
*Dummy Scans*: 4

The table summarizes the used conditions for scans:

| Pulse Width | **$10^{o}$** |
|---|---|
| | RT |
| Relaxation delay | 0 |
| Acquisition time | 4s |
| Spectral Window | 30 ppm |
| Transmitter Offset | 7.5 ppm |
| Number of Scans for 400 MHz | 512 |

(3) Post-acquisition processing

Apodization: LB = 0.1 Hz
Zero-filling: 256 K
Phasing: Manual phase correction
Baseline correction: Bernstein Polynomials

(4) Quantitative measurement of integrals

The proton integrals and range (ppm) of all signals used for quantification have been documented below. The integral of the internal calibrant resonance signal (dimethyl sulfone) is in bold.

| **1** | |
|---|---|
| Range | Absolute |
| 7.06-7.00 | 1.02 |
| 6.98-6.93 | 1.01 |
| 3.77-3.70 | 2.03 |
| 3.22-3.16 | 2.00 |
| **2.99-2.94** | **6.08** |
| $\boldsymbol{Int_t}$ = 1.01, $\boldsymbol{n_t}$ = 1<br>$\boldsymbol{MW_t}$ = 325.32 g/mol | |

| **2** | |
|---|---|
| Range | Absolute |
| 7.30-7.26 | 0.99 |
| 3.14-3.08 | 1.05 |
| 3.69-3.63 | 2.00 |
| 3.21-3.15 | 2.00 |
| **3.00-2.93** | **37.09** |
| $\boldsymbol{Int_t}$ = 1.01, $\boldsymbol{n_t}$ = 1<br>$\boldsymbol{MW_t}$ = 348.33 g/mol | |

| **3** | |
|---|---|
| Range | Absolute |
| 7.35-7.30 | 1.01 |
| 7.04-7.00 | 1.01 |
| 3.71-3.65 | 2.00 |
| 3.25-3.18 | 2.05 |
| **3.00-2.95** | **15.51** |
| $\boldsymbol{Int_t}$ = 1.01, $\boldsymbol{n_t}$ = 1<br>$\boldsymbol{MW_t}$ = 373.34 g/mol | |

| **4** | |
|---|---|
| Range | Absolute |
| 7.35-7.31 | 1.00 |
| 7.06-6.98 | 1.99 |
| 3.70-3.62 | 2.00 |
| 3.20-3.13 | 2.21 |
| **3.01-2.94** | **45.48** |
| $\boldsymbol{Int_t}$ = 1.02, $\boldsymbol{n_t}$ = 1<br>$\boldsymbol{MW_t}$ = 382.77 g/mol | |

| **5** | |
|---|---|
| Range | Absolute |
| 7.19-7.13 | 1.01 |
| 7.03-6.97 | 1.02 |
| 3.69-3.62 | 2.00 |
| 3.19-3.13 | 2.09 |
| **3.00-2.95** | **22.52** |
| $Int_t$ = 1.02, $n_t$ = 1<br>$MW_t$ = 426.02 g/mol | |

(5) Calculation

$$P\% = \frac{n_{IC} \times Int_t \times MW_t \times m_{IC}}{n_t \times Int_{IC} \times MW_{IC} \times m_s} \times P_{IC}$$

- $P$ – purity of the target analyte, %
- $P_{IC}$ – purity of the internal calibrant, %
- $n_{IC}$ – number of protons that give rise to $Int_{IC}$
- $n_t$ – number of protons that give rise to $Int_t$
- $Int_{IC}$ – integral of the internal calibrant resonance signal
- $Int_t$ – integral of the target analyte resonance signal
- $MW_{IC}$ – molecular weight of the internal calibrant
- $MW_t$ – molecular weight of the target analyte
- $m_{IC}$ – weight of the internal calibrant
- $m_s$ – weight of the target analyte

| Compounds ID | Purity (%) |
|---|---|
| **1** | 99.71 |
| **2** | 99.36 |
| **3** | 99.75 |
| **4** | 99.96 |
| **5** | 99.80 |

### Uncropped images for SI Fig. 8

Replicate #1

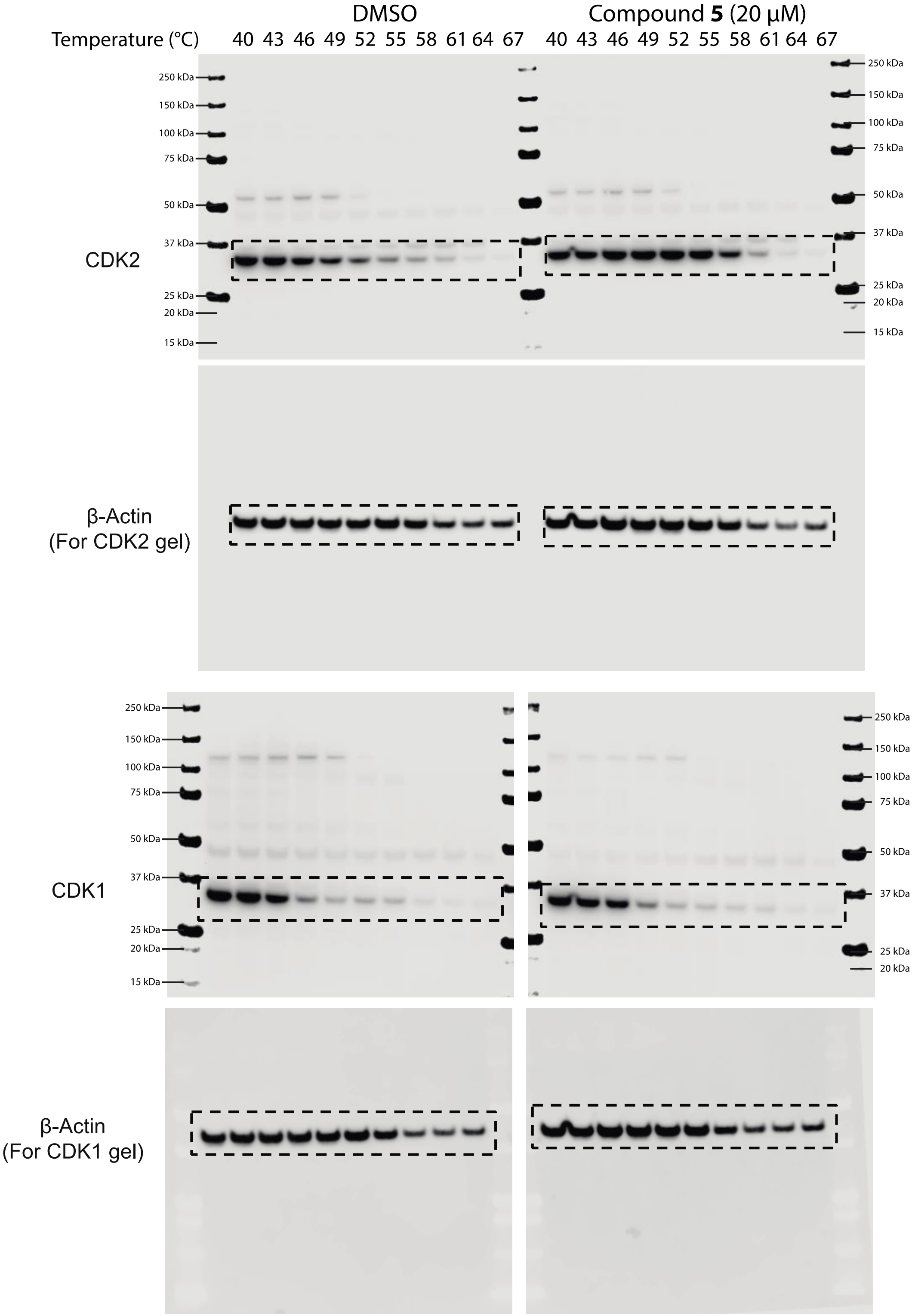

### Uncropped images for SI Fig. 8 Replicate #2

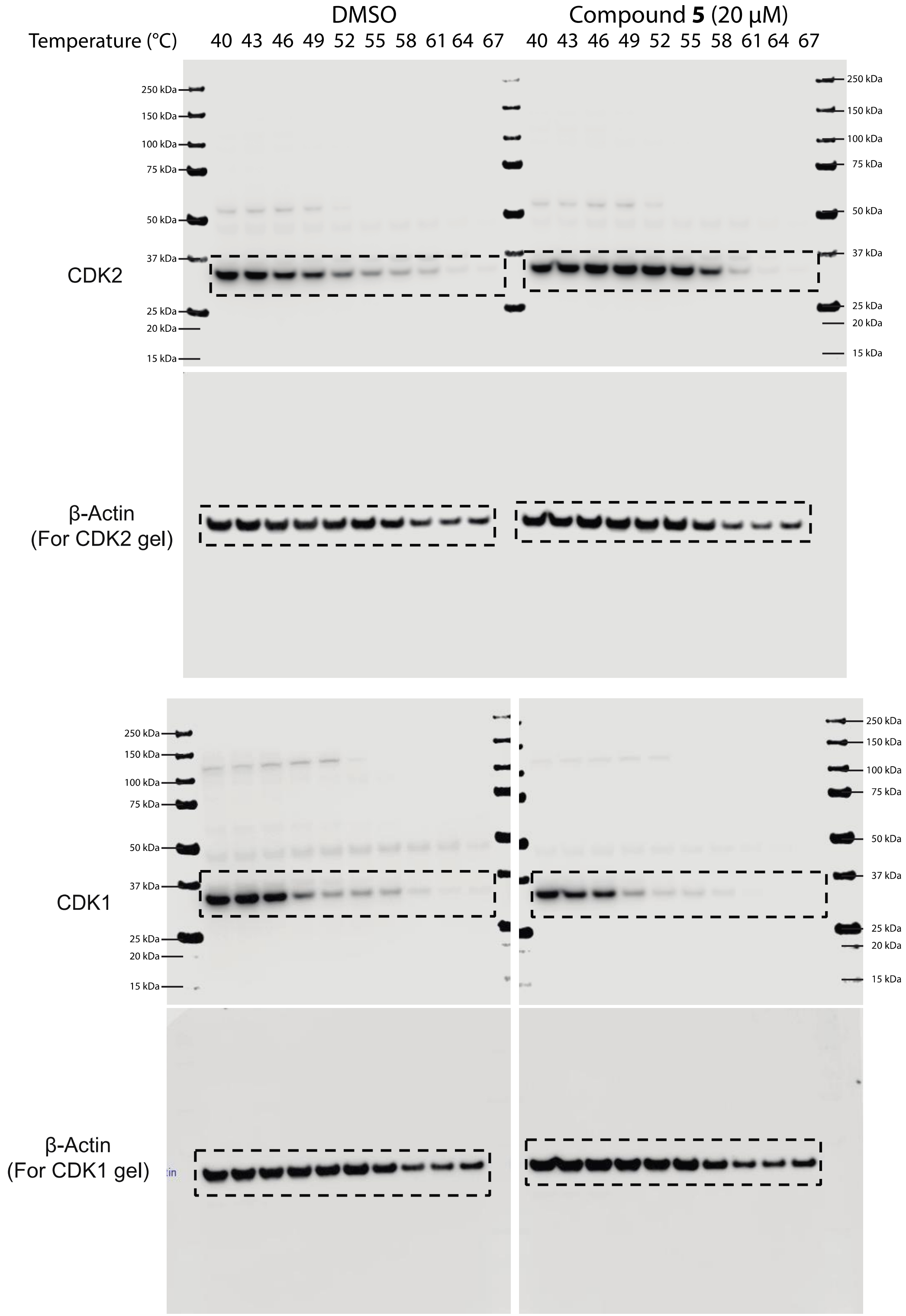

**Uncropped images for SI Fig. 8** Replicate #3

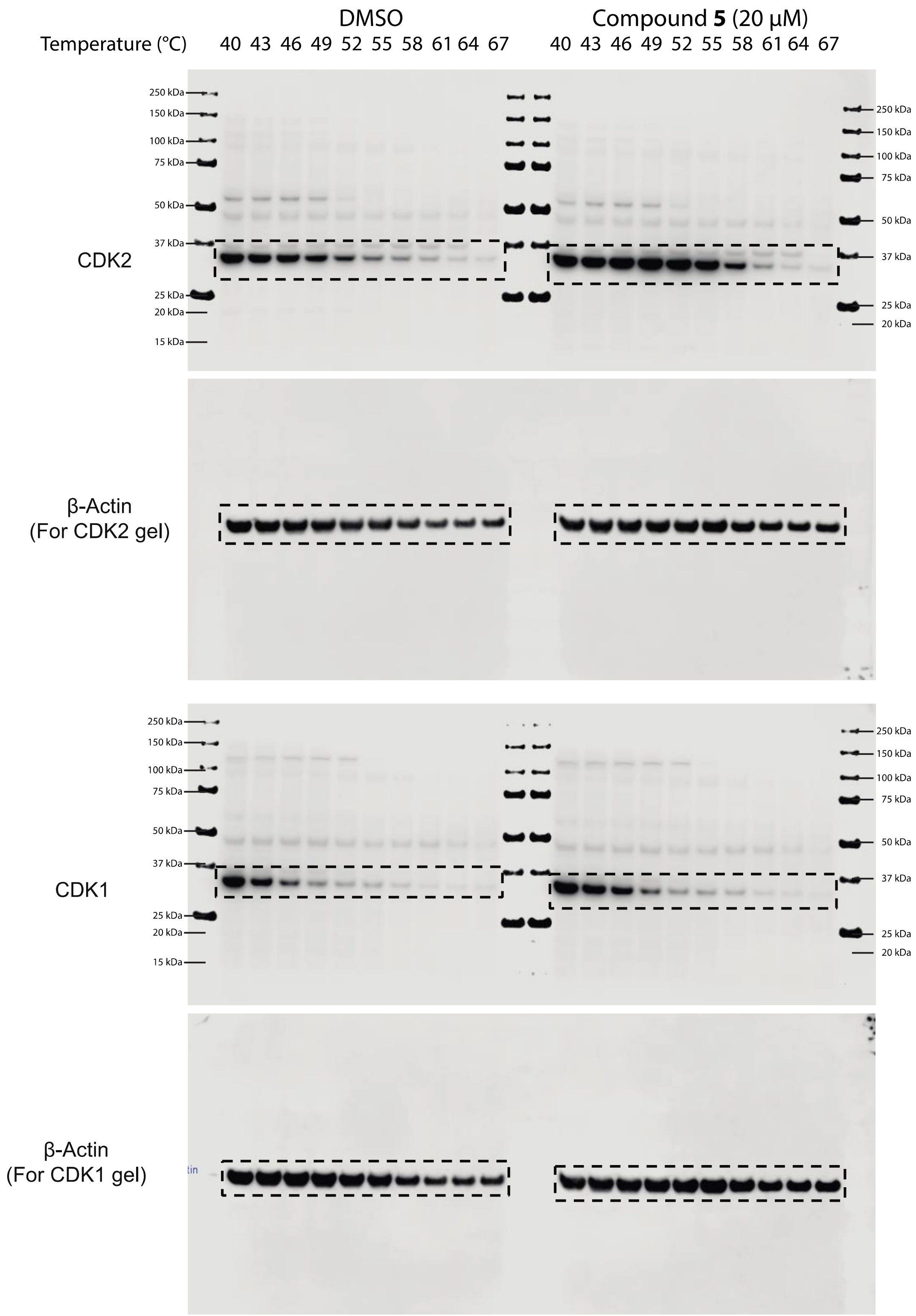

### Uncropped images for SI Fig. 11 Replicate #1

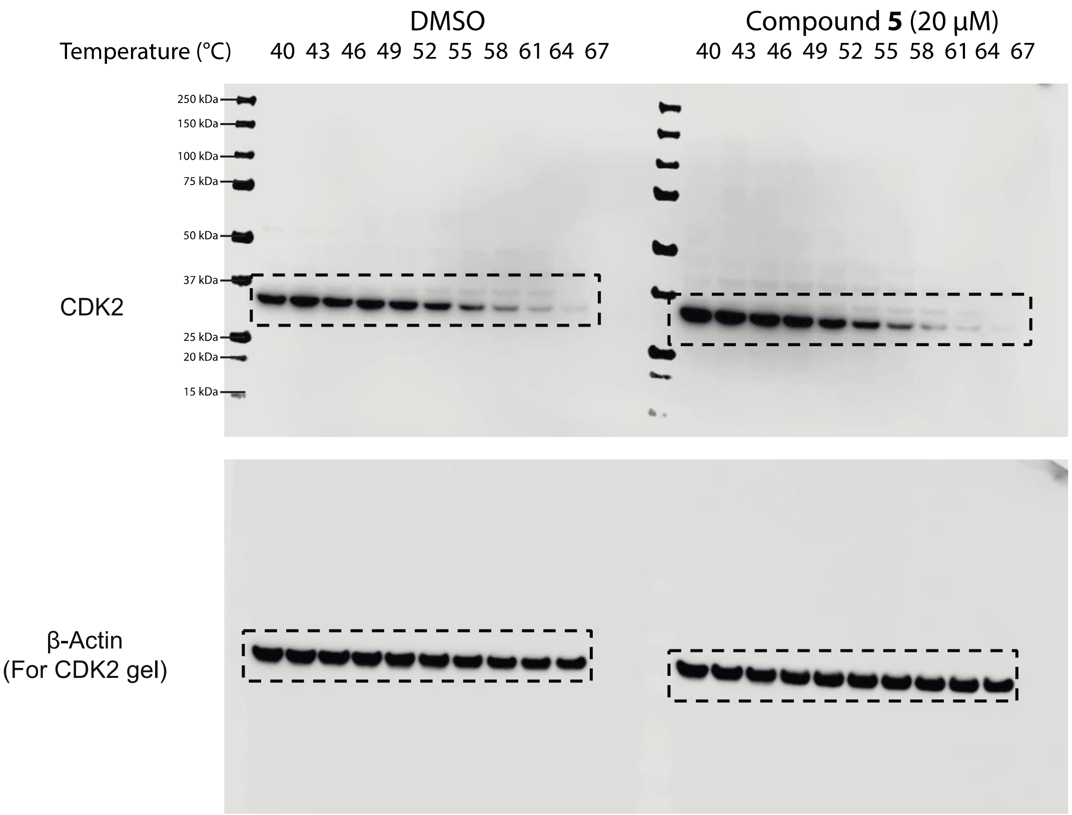

### Uncropped images for SI Fig. 11 Replicate #2

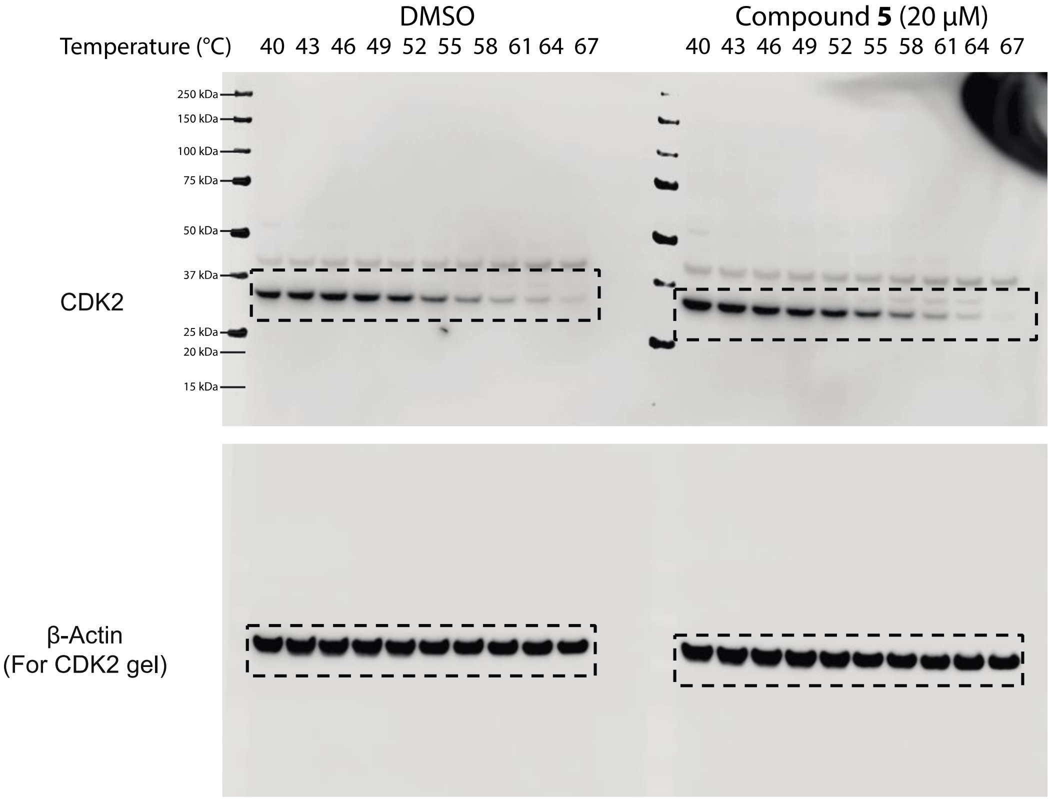

### Uncropped images for SI Fig. 11 Replicate #3

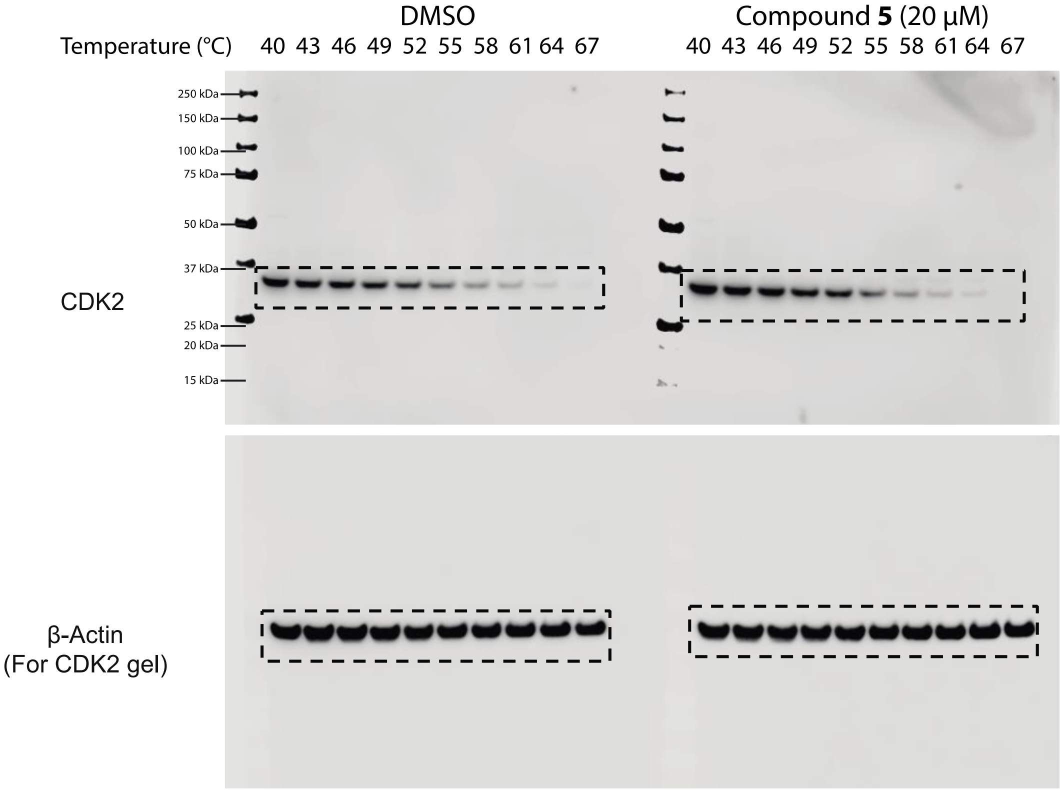
